# Evaluating proteome allocation of *Saccharomyces cerevisiae* phenotypes with resource balance analysis

**DOI:** 10.1101/2022.09.20.508694

**Authors:** Hoang V. Dinh, Costas D. Maranas

## Abstract

*Saccharomyces cerevisiae* is an important model organism and a workhorse in bioproduction. Here, we reconstructed a compact and tractable genome-scale resource balance analysis (RBA) model (i.e., *sc*RBA) to analyze metabolic fluxes and proteome allocation in a computationally efficient manner. Resource capacity models such as *sc*RBA provide the quantitative means to identify bottlenecks in biosynthetic pathways due to enzyme, compartment size, and/or ribosome availability limitations. ATP maintenance rate and *in vivo* apparent turnover numbers (k_app_) were regressed from metabolic flux and protein concentration data to capture observed physiological growth yield and proteome efficiency and allocation, respectively. Estimated parameter values were found to vary with oxygen and nutrient availability. Overall, this work (i) provides condition-specific model parameters to recapitulate phenotypes corresponding to different extracellular environments, (ii) alludes to the enhancing effect of substrate channeling and post-translational activation on *in vivo* enzyme efficiency in glycolysis and electron transport chain, and (iii) reveals that the Crabtree effect is underpinned by specific limitations in mitochondrial proteome capacity and secondarily ribosome availability rather than overall proteome capacity.

## Introduction

*Saccharomyces cerevisiae* is a prototrophic yeast (unicellular fungus organism) that has been domesticated for making bread and wine since ancient times^1^. It occupies a wide spectrum of natural habitats, ranging from soils, insects, grapes, and leaves and trunks of plant species^1^. The organism is well-known for its ethanol production (and tolerance) even under aerobic conditions (i.e., the Crabtree effect)^2^. *S. cerevisiae* have been considered as a “generally recognized as safe” (GRAS) organism and used extensively in biological research as a model eukaryotic organism^3^ and in large-scale fermentation^4^. A prime example is its use in producing bioethanol^5^ for which global demand was 28.91 billion gallons in 2019^6^. *S. cerevisiae* has also been extensively re-engineered by metabolic engineering to produce various compounds such as fatty acid derivatives and biofuels^7^, building block organic acids^8^, biopharmaceutical proteins^9^, natural products^10,11^, and food additives^12^. For example, industrial-scale production of the antimalarial drug artemisinin’s precursors has been achieved using yeast strain with heterologous expression of *Artemisia annua*’s enzymes^13^. These numerous applications and adaptations stem from the organism’s robustness in industrial settings (e.g., resistance to growth inhibitors, pH, osmotic, and ethanol stresses)^14^ and a strong engineering foundation established by well-characterized genome sequences and annotations^15,16^ and availability of synthetic biology tools^17^.

These has been a number of genome-wide systems biology studies for *S. cerevisiae* focusing on cellular expression under growth condition perturbations^18^, regulation^19,20^, metabolism^21^, and genotype-phenotype correlation^22^. The study of *S. cerevisiae* metabolism has received considerable attention. Starting from the first genome-scale metabolic (GSM) model reconstructed in 2003^23^, successive models have been developed with improved coverage of genome and metabolic functions^24,25^. Using as inputs only biomass composition, gene annotations, gene to protein to reaction (GPR) associations, and reaction reversibility information, GSM models have been shown to predict metabolic fluxes and theoretical yields reasonably well^21^. They have been used extensively to suggest genetic perturbation strategies for metabolic engineering^24^. Recently, upgrades of GSM models accounting for protein and enzyme availability limitations have been made to improve model prediction by imposing upper limits to metabolic fluxes. Sánchez and coworkers developed the GECKO framework that imposes flux upper bounds derived from protein concentration measurements^26^. Oftadeh and coworkers presented the expression and thermodynamics flux (ETFL) model that account for cellular expression system and reaction thermodynamics^27^. These models display enhanced predictions for batch culture conditions where nutrients are in excess and enzyme production capacity becomes the bottleneck. Beyond metabolism and expression processes, a whole-cell model containing 26 sub-models within has also been developed by Ye and coworkers to capture holistically cellular processes^28^.

In this paper, we put forth a computationally tractable to parameterize and simulate resource balance analysis (RBA) model for *S. cerevisiae* referred to as *sc*RBA that focuses on metabolism and enzyme and ribosome production. The goal is to construct a model with a level of detail that experimental data can support parameter estimation. Sets of growth and non-growth associated ATP maintenance parameters specific to growth conditions are regressed from a large collection of *S. cerevisiae* growth phenotype data^29–48^ to accurately predict condition-dependent growth yield. The inferred ATP maintenance rates increase significantly in the presence of oxygen and under carbon and nitrogen limitation in agreement with known yeast physiology^49–56^. *In vivo* enzyme turnover parameters (k_app_) (indicating enzyme efficiency) were regressed using multiple measured extracellular fluxes and protein concentration datasets^29,30,34,37^. We found that k_app_ values are often very different than catalogued *in vitro* turnover numbers (k_cat_)^57^ which are typically used in proteome allocation metabolic models. Notably for 4 out of 10 enzymes in glycolysis, 4 out of 4 in electron transport chain, ATP synthase, and 10 out of 20 in amino acyl-tRNA synthetase pathways estimated k_app_ values were significantly larger (i.e., by up to 189-fold for hexokinase) than tabulated k_cat_ data reflecting higher *in vivo* enzymatic efficiency. While insight into the exact mechanism for this enhancement is not revealed by the parameterization of the RBA model, there is ample literature evidence^58–64^ for the presence and function of *in vivo* metabolons and enzyme post-translational activations. Interestingly, inferred k_app_ values were generally lower for (i) alternate carbon substrates, (ii) anaerobic conditions, or (iii) carbon/nitrogen limitations in accordance with lower enzymatic efficiencies implied by the phenotypic data. The parameterized *sc*RBA model for *S. cerevisiae* without any tuning quantitatively predicted ethanol overflow under abundant glucose and oxygen conditions. Mitochondrial proteome capacity and secondarily ribosome availability rather than total proteome capacity were revealed as the limiting resources for faster growth utilizing ATP from respiration. Ethanol overflow provides an alternative redox-balanced mechanism to produce ATP for growth on glucose leveraging remaining rRNA and non-mitochondrial proteome capacity. The *sc*RBA model captured the flux-limiting effect of enzyme and/or ribosomes availability (i.e., as low as below 20% of FBA predicted fluxes) for 78% of metabolic reactions under glucose uptake conditions by disallowing ATP-wasting cycles that require unavailable proteome resources. *sc*RBA based predicted maximal product yields for 28 biochemicals were generally only marginally reduced compared to FBA-calculated values. This is because maximum production pathways are unlikely to contain futile cycles. Exceptions to this include succinate whose RBA-based maximum theoretical yield was only 21% of the FBA-predicted yield due to the enzymatically inefficient pyruvate-to-succinate conversion steps in yeasts. Overall, this work puts forth the *sc*RBA model for *S. cerevisiae* that draws from condition-specific parameters to improve prediction accuracy. Its applicability is demonstrated through case studies of *S. cerevisiae*’s metabolic flux and product yield limit estimations.

## Methods

### RBA model reconstruction

The model *sc*RBA consists of (macro)molecules and reactions for metabolism and cellular machinery production linked through steady-state mass balance constraints as in FBA^65^ (see Fig. 1a for a schematic representation). Here, we provide data sources necessary for reconstruction and briefly explain the model elements linking (macro)molecules and reactions. The reconstruction method is described in detail in the Supplementary Methods, with user instructions, formulation (adapted from Goelzer et al., 2011^66^), and indexing. Metabolites and metabolic reactions are ported from *iSace*1144 (available at https://github.com/maranasgroup/iSace_GSM). Blocked reactions identified by flux variability analysis^67^ were excluded from *sc*RBA. Protein translation reactions entail amino acids (in the form of charged-tRNA), cofactors, and energy in the form of GTP and ATP^68,69^ with stoichiometric coefficients designed to match the corresponding protein sequences^16^ using the Uniprot database^70^. The mass action contribution of ribosomes is set to be proportional to the sum of all protein translation fluxes. Ribosomes, in turn, are synthesized from proteins and rRNAs. In yeast, most of the genes (i.e., 1,201 out of 1,208 genes in the model) are encoded in the nucleus chromosome and translated by the nuclear ribosome. The remaining mitochondrial genes^71^ (i.e., 7 genes in the model) are translated using reactions that utilize the mitochondrial ribosome which is treated as separate from the cytosolic counterpart. The stoichiometric coefficients for the proteins and rRNAs associated with ribosome production are obtained from the SGD database^16^ and ribosome structure observations^72^. rRNA relative ratios are sourced from the RNAcentral database^73^. Enzymes are formed from the corresponding protein subunits whose stoichiometric coefficients are obtained from the Uniprot database^70^. Biomass precursor producing reactions are included in *sc*RBA to inventory all enzymes, ribosomes, and all other macromolecules needed to form biomass. Precursors (and their relative compositions) of DNA, lipids, carbohydrates, metal ions, sulphate, phosphate, and cofactors are ported from the GSM model *iSace*1144. Based on experimental macromolecular measurements^29,38,74,75^ the mass fractions of most macromolecules are assumed to remain invariant except for protein, RNA, and carbohydrate fractions that change with increasing growth rate (see Supplementary Data 1). Instead of reconstructing multiple models with different biomass coefficients at different growth rates, the biomass reaction is recast as a set of precursor sink reactions whose fluxes are equal to the coefficients multiplied by the growth rate. We ensure that the biomass molecular weight is always 1 g mmol^-1^ so that growth yield and rate predictions are consistent^76^. Detailed biomass formulation is provided in the Supplementary Methods and Supplementary Data 1.

**Fig 1.**
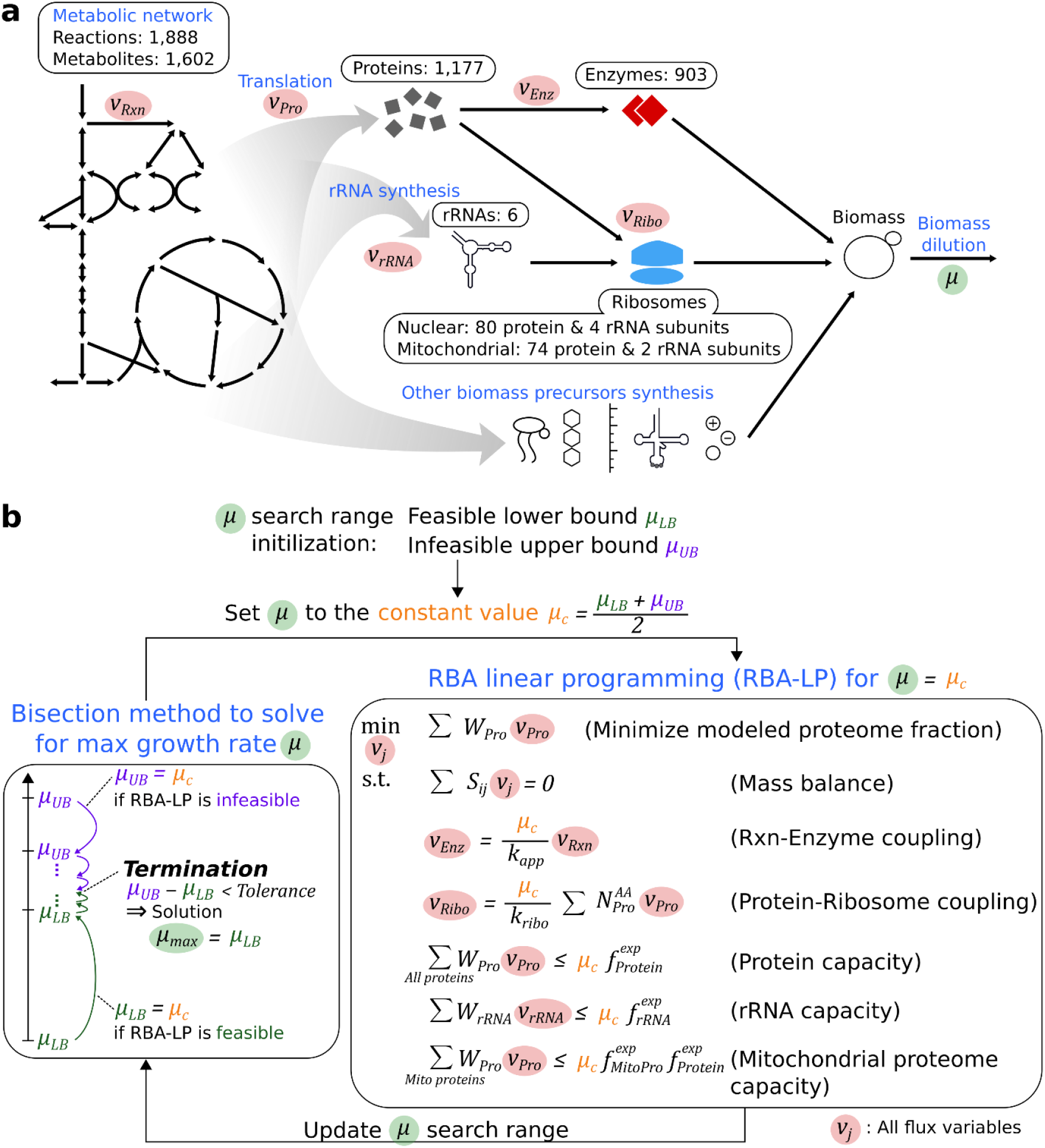
(**a**) Schematic representation of the *sc*RBA model (macro)molecular participants and reactions. (**b**) Overview of bisection method and the RBA linear programming (RBA-LP) formulation that are solved iteratively to obtain the maximal growth rate. Flux variables are highlighted in red and the growth rate variable is highlighted in green. The topology of all *sc*RBA model captured variables are shown in Fig. 1a. Model parameters are briefly explained in the text and formulation details are available in the Supplementary Methods.

The total amount of protein, enzyme, and ribosome produced is determined by the reaction-enzyme and protein-ribosome coupling constraints and limited by the protein and rRNA capacity constraints (see Fig. 1b for a formulation overview, Supplementary Methods for the complete formulation details, and Goelzer et al., 2011^66^ for derivations). The k_app_ parameter values in the reaction-enzyme coupling constraint are derived from experimental flux and proteomics data (see “Estimation of *in vivo* k_app_” in Methods). In the protein-ribosome coupling constraint, the protein sequence length values 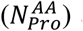 are taken from the SGD database^16^ and the ribosome efficiency parameter (k_ribo_) is fitted using growth phenotype data^30^. From the experimentally derived^77^ average value of 10.5 amino acids elongated per ribosome per second, we re-parameterized the k_ribo_ value by successively increasing it from 10.5 in increments of 0.1 until the predicted growth rate matched the highest reported experimental value of 0.49 h^−1^ (in rich media)^30^. This was met for a slightly higher value of k_ribo_ of 13.2 amino acids per ribosome per second. In the *sc*RBA model, enzyme and ribosome production is limited by the experimentally measured protein and rRNA levels^29,38,74,75^ through capacity constraints. Molecular weights of protein and rRNA (i.e., *W*_*Pro*_ and *W*_*rRNA*_) are used to convert molar amounts to grams which are set to less than the experimentally found limits (i.e., 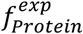 and 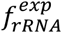) in the capacity constraints. Production of proteins locating in mitochondria are also limited to a mass fraction (i.e., 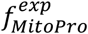) of the total proteome capacity. For glucose uptake conditions the reported mitochondria per cell volume/volume ratio of 5% ^78^ is used. This 5% threshold for mitochondrial proteins could be mechanistically explained as reflecting membrane surface area and inner volume limits for mitochondrial metabolic pathways (e.g., TCA cycle and electron transport chain) and ribosomes. *sc*RBA directly accounts for only proteins participating in metabolism and translation/elongation. Proteins involved in other processes such as protein folding chaperone and cellular maintenance are not functionally part of *sc*RBA. We assumed that the modeled proteome including metabolic and ribosomal proteins is 55% of the total proteome^27^. The cost of producing the remaining 45% is accounted for in an aggregate manner assuming average amino acid composition^74^. In resource allocation models this is formulated by adding a reaction producing a non-functional so-called “dummy” protein^27,79^. The model accounts explicitly the six rRNA species that are part of ribosomes (see Supplementary Methods for details) which constitute as much as 80% of total RNA^80^. For computational efficiency, mRNA and tRNA demand (i.e., reserving 5% and 15% of total RNA, respectively) are accounted for in an aggregate manner assuming average composition^23^. Similar to the proteome, a reaction producing a non-functional RNA is added to the model. *S. cerevisiae* maintains reserved ribosome^77^ and enzyme^35^ capacity which makes up the difference between experimentally observed and required amounts (estimated under nitrogen limited conditions^35,77^). The reserved proteome capacity is maintained to prepare cells for changes in growth conditions in the same manner that reserved ribosome capacity enables faster growth immediately upon nutrient availability upshift^35,77^. In model, the amounts of non-functional protein and RNA representing reserved capacity are treated as fitted variables so as to recapitulate reserved capacity being present or exhausted depending on growth conditions.

### Estimation of ATP maintenance rates

ATP maintenance rates are parameters used in both FBA and RBA model to account for the energy cost of replicating cells. Growth-associated ATP maintenance (GAM) (in mmol gDW^−1^) rate captures the energy demand per unit of produced biomass. Non-growth associated ATP maintenance (NGAM) (in mmol gDW^−1^ h^−1^) rate captures the energy demand associated with cellular processes such as repair and maintenance^81^. GAM_FBA_ (i.e., GAM in FBA model) and NGAM parameters were regressed from growth phenotype datasets recorded at different growth rates using the FBA model *iSace*1144. For every dataset, ATP maintenance rate was estimated by maximizing flux through the ATP hydrolysis reaction (i.e., *ATP* + *H*_2_*O* → *ADP* + *P*_*i*_ + *H*^+^) subject to experimentally measured extracellular fluxes and growth rate^29–37,41–48^. NGAM parameter was equal to the maximal ATP hydrolysis flux estimated from growth-arrested data^38–40^. GAM_FBA_ is the slope of a linear regression of maximal ATP hydrolysis flux vs. growth rate values whereas the intercept is the NGAM value. GAM_RBA_ (i.e., GAM in RBA model) is estimated by subtracting from GAM_FBA_ value the portion equivalent to protein translation elongation’s energy cost, which is approximately 2 mmol of ATP per mmol of amino acid^27^. The subtracted amount is 7.6 mmol ATP gDW^−1^, derived from experimental amino acid measurements^74^. The NGAM parameter is estimated from growth-arrested data where neither biomass nor protein synthesis is underway and thus the parameter is the same for both FBA and RBA. Different GAM_FBA_ and NGAM parameter sets were regressed from datasets under the following growth conditions: (i) (nutrient-abundant) batch and anaerobic or microaerobic, (ii) C-limited chemostats and anaerobic or microaerobic, (iii) batch or C-limited chemostats and aerobic, (iv) N-limited chemostats and aerobic. Experimental flux inputs and calculated results are recorded in Supplementary Data 2.

### Estimation of *in vivo* k_app_

k_app_ was calculated by dividing estimated intracellular metabolic fluxes by experimental enzyme concentrations^82^ (see Supplementary Methods for the workflow and Supplementary Data 3 for details). From literature-reported data^29,30,34,37^, different k_app_ parameter sets were determined for growth conditions under (i) (nutrient-abundant) batch/glucose, (ii) batch/galactose, (iii) batch/maltose, (iv) batch/trehalose, (v) C-limited chemostats/glucose D = 0.1 h^−1^ and (vi) D = 0.3 h^−1^, and (vii) N-limited chemostats/glucose D = 0.1 h^−1^.

### Growth maximization, flux variability analysis, and predicted yield using *sc*RBA and FBA

An overview of the RBA optimization model that identifies the maximal growth rate is provided in Fig. 1b. By fixing the growth rate the *sc*RBA model is converted into a linear programming formulation (i.e., RBA-LP) which can be efficiently solved. An iterative method is employed to converge the upper (infeasible) and lower (feasible) bounds on growth rate within a tolerance criterion of 10^−5^ h^−1^.

In analogy to flux variability analysis (FVA)^67^, lower and upper bounds of reaction fluxes can be calculated using *sc*RBA and FBA models by updating the objective function of the model (see Fig. 1b) to the minimization or maximization of the flux in question and imposing the experimental glucose uptake rate of 13.2 mmol gDW^−1^ h^−1^ and growth rate of 0.42 h^−1 30^. Experimental (absolute) glucose uptake and growth rates were used in the simulations to be consistent with model parameters derived from absolute flux and concentration measurements. Flux ranges under FBA and RBA are contrasted to elucidate the role of capacity constraints on the flux allocation flexibility. Maximal compound production rate was identified by maximizing the corresponding (sink) exchange reaction flux variable subject to a glucose uptake of 13.2 mmol gDW^−1^ h^−1 30^ and growth rate set at a minimum of 0.1 h^−1^. Hexadecanoic acid and hexadecanol (i.e., C_16_) were used as proxies in model for *in vivo* mixture of free fatty acids and fatty alcohols of different chain lengths, respectively. Heterologous pathways for the synthesis of tested compounds were reconstructed based on previous studies^7,83–109^, as necessary. The RBA-predicted maximal production yield (i.e., 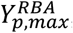, in g g-Glucose^−1^) was calculated using the following equation:

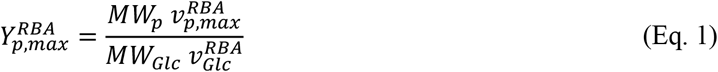

where 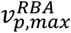 is the RBA-predicted maximal production rate, *MW*_*p*_ is the product molecular weight, ν_*Glc*_ is the RBA-predicted glucose uptake rate, and *MW*_*Glc*_ is the molecular weight of glucose. FBA-calculated maximal production yield (i.e., 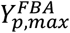) was determined using Eq. 1 but with FBA-predicted rather than RBA-predicted quantities. Fluxes are in mmol gDW^−1^ h^−1^ and molecular weights are in g mmol^−1^.

### Software implementation

COBRApy^110^ with IBM ILOG CPLEX solver (version 12.10.0.0) were used for FBA model optimization. The FBA model *iSace*1144 is available at https://github.com/maranasgroup/iSace_GSM^111^. General Algebraic Modeling System (GAMS) programming language (version 39.1.0, GAMS Development Corporation) with Soplex solver (version 6.0)^112^ was used for RBA model k_app_ parameterization and optimization. Input files as excel spreadsheets were used to build the RBA model in GAMS format. Python 3.6 was used as the central platform to run all mentioned processes. All scripts and input and output files are available in the GitHub repository https://github.com/maranasgroup/scRBA.

## Results

### Estimation of growth and non-growth associated ATP maintenance

ATP maintenance rate parameters, GAM and NGAM are necessary to accurately account for the fraction of glucose uptake apportioned towards energy production and ultimately growth^76^.We estimated GAM and NGAM parameters values using an ATP synthase proton/ATP ratio of 10/3. This reflects the fact that the *S. cerevisiae* ATP synthase consists of 10 c-ring for every 3 F_1_F_0_ subunits^113^ resulting in 10 proton molecules translocated across mitochondrial membrane per 3 ATP molecules produced. Note that earlier GSM models used a proton/ATP ratio of 12/3 ^23,114^. An overview of the literature-reported experimental datasets, methods, and GAM/NGAM values is provided in Fig. 2. We calculated from metabolic fluxes of growth-arrested cells^38–40^ the NGAM values (in mmol gDW^−1^ h^−1^) for (i) aerobic C-limited (NGAM = 1.0), (ii) anaerobic C-limited (NGAM = 1.0), and (iii) aerobic N-limited (NGAM = 3.9). Obtained results reveal a largely unchanging NGAM value across C-limited aerobic and anaerobic conditions even though earlier studies pointed at some small differences^39^. New results are possibly due to the updated ATP synthase reaction stoichiometry reflecting recent literature information^113^. Note that the calculated NGAM value is nearly 4-fold higher under nitrogen-limited conditions indicating more energy is expended towards cellular maintenance due to the limited nitrogen pool.

**Fig. 2.**
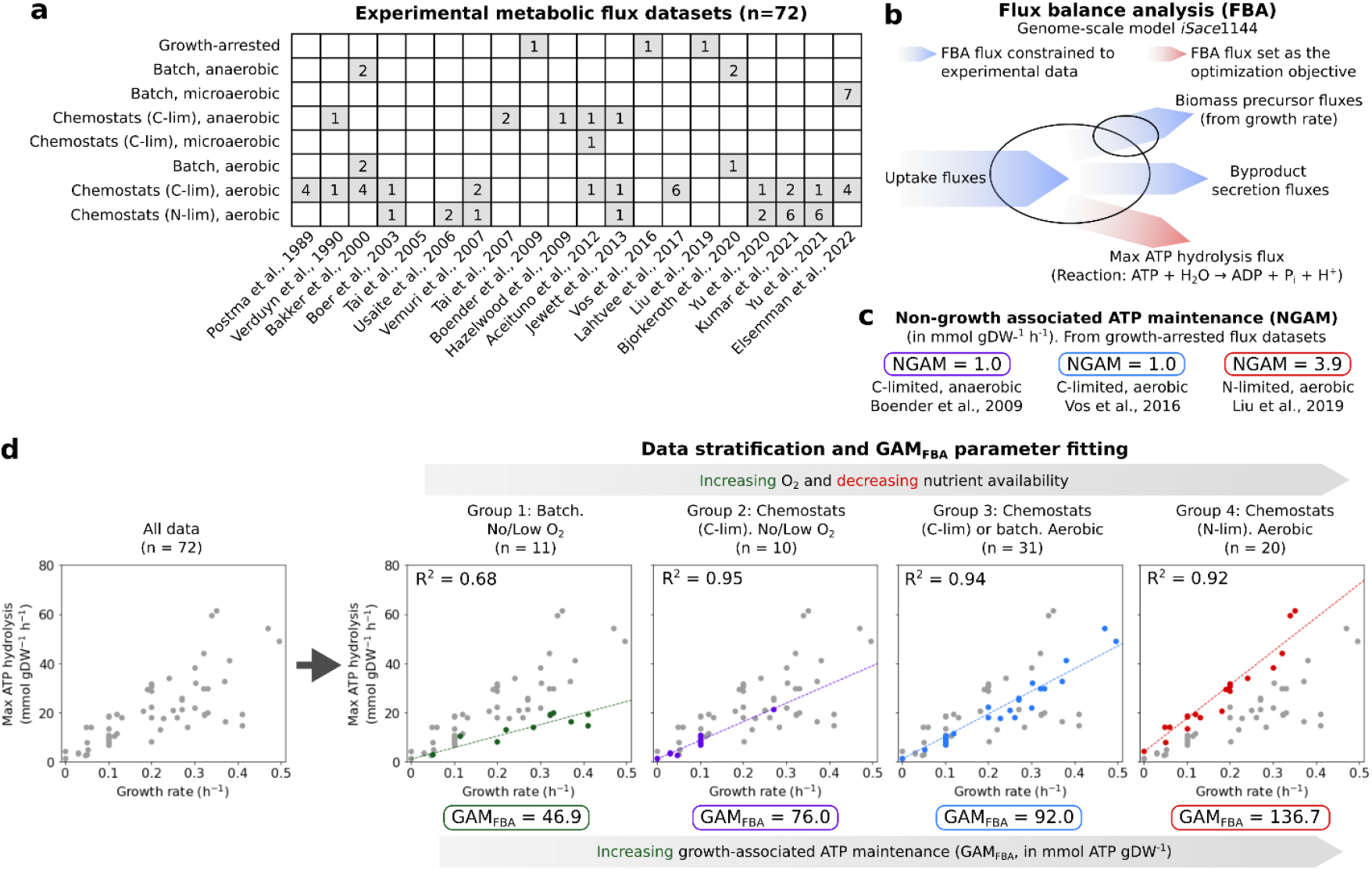
Parameterization of growth and non-growth associated ATP maintenance. (**a**) Summary of the experimental datasets. (**b**) Overview of the FBA procedure to calculate maximal ATP hydrolysis rate. (**c**) Non-growth associated ATP maintenance (NGAM) values. (**d**) Growth-associated ATP maintenance (GAM) values. R^2^ is the coefficient of determination for the linear regression to determine the GAM value as the slope. Overall, GAM and NGAM values increase under nutrient limitation and in the presence of oxygen.

GAM_FBA_ values (in mmol gDW^−1^) are regressed from metabolic flux datasets at different growth rates. Significantly different GAM_FBA_ values were inferred for subsets of data under different experimental oxygen and nutrient availability (see Fig. 2d). This means that GAM_FBA_ values must be tailored to the growth conditions to maintain fidelity of prediction. The lowest GAM value (i.e., GAM_FBA_ = 46.9) is associated with anaerobic batch conditions where nutrients are abundant. Carbon limitation (i.e., GAM_FBA_ = 76.0), presence of oxygen (i.e., GAM_FBA_ = 92.0), and nitrogen limitation (i.e., GAM_FBA_ = 136.7) increase GAM_FBA_ by 1.6-fold, 2-fold, and 2.8-fold, respectively. In the presence of oxygen, redox energy is partly diverted towards dissipating reactive oxygen species from respiration^49,50^. Respiration also requires functional mitochondria whose synthesis and damage repair require energy input^51,52^. The lower GAM_FBA_ value under anaerobic conditions may also be in part due to *S. cerevisiae* adaptation to conditions with abundant fruit-borne sugar but low oxygen availability due to diffusion limitations^115^. Under glucose-limitation, higher ATP maintenance rate is likely needed for the synthesis of the catabolic catalytic apparatus to scavenge alternate carbon sources other than glucose (e.g., through the Snf1 regulatory pathway)^53^. While this is an important adaptation in the natural habitat, it is counterproductive in laboratory settings where alternative carbon sources are not provided. Under N-limitation, higher ATP maintenance rate is associated with the degradation of selected proteins (e.g., ribosomal proteins)^54,55^ to replenish depleted pools of amino acids and other N-containing compounds^56^. Because nitrogenous metabolite pools are not conserved by downregulating protein synthesis but rather by engaging an ATP-consuming protein synthesis-degradation cycle, this leads to a significantly higher GAM_FBA_ value under N-limitation. Overall, we estimated condition-specific GAM and NGAM values and provided hypotheses for mechanistic bases for *S. cerevisiae*’s varied ATP maintenance under different growth conditions. Interestingly, we found that under P-limitation^36^ ATP maintenance does not follow a linear trend (as shown in Fig. 2d) and GAM value (i.e., slope) becomes dependent on the degree of limitation (see Supplementary Fig. 1).

### Estimation of *in vivo* apparent k_app_ values

Enzyme turnover numbers k_app_ are RBA model parameters that directly affect proteome allocation needs as their product with enzyme levels must exactly match the metabolic flux values. Underestimated values for k_app_ result in higher than needed requirements for protein levels and vice-versa. Even though *in vitro* turnover numbers (k_cat_) for many reactions are available in the literature and biochemical databases such as BRENDA^57^, they are not immediately usable in RBA models. This is because k_cat_ entries do not capture the nuances of the *in vivo* environment (e.g., sub-saturation of enzyme, substrate channeling, post-translational modifications, etc.) that can dramatically alter enzymatic efficiencies^82,116^. In model *sc*RBA, we instead rely on k_app_ values supported by measured metabolic fluxes and corresponding enzyme levels. Assuming that the *in vivo* substrate concentration is below the saturated level, k_app_ values are by definition lower than k_cat_ values^57^ (i.e., in the Michaelis-Menten expression, *k*_*app*_ = *k*_*cat*_([*S*]/(*K*_*M*_ + [*S*])) ≤ *k*_*cat*_). However, we found in agreement with earlier studies^82,117^ that the reverse is true for several enzymes (i.e., 46 out of 132 enzymes in *E. coli* based on available data^82^) indicating that enzyme efficiency is often enhanced *in vivo* through possibly substrate channeling and/or enzyme activation (see Fig. 3 for a visual and Supplementary Data 4 for tabulated values). Often the enzyme enhancement outpaces substrate saturation effects. Enzymes with experimental evidence confirming *in vivo* activity enhancement are reported in Table 1. Enzymes without direct evidence but with predicted marked enhancements *in vivo* are reported in Table 2 as candidates for further testing. Among these candidates is included a hypothetical model of a metabolon involving the ATP synthase proposed for yeast in analogy to the mammalian counterpart^118^. We also found that k_app_ values are lower than k_cat_ for four enzymes in the glycolysis metabolon^58^ (see Fig. 3) which suggests that the reducing effect of enzyme sub-saturation is stronger than any enhancing effect of substrate channeling. We find that *in vivo* enhancements occur predominantly in high-flux glycolysis, electron transport chain, and ethanol fermentation alluding to a mechanism to reduce proteome investment for high-capacity enzymes. Note that proteome allocation needs towards these pathways would have been significantly higher if the *in vivo* k_cat_ values were unaltered from the *in vitro* ones. This makes sense as pathways with high flux would be under a stronger selection to achieve *in vivo* k_app_ enhancements through a variety of mechanisms in comparison to low flux routes. This result further reinforces the need to estimate and utilize condition-specific *in vivo* k_app_ values to faithfully recapitulate *in vivo* enzymatic efficiency and predict proteome allocation with RBA models.

**Table 1.**
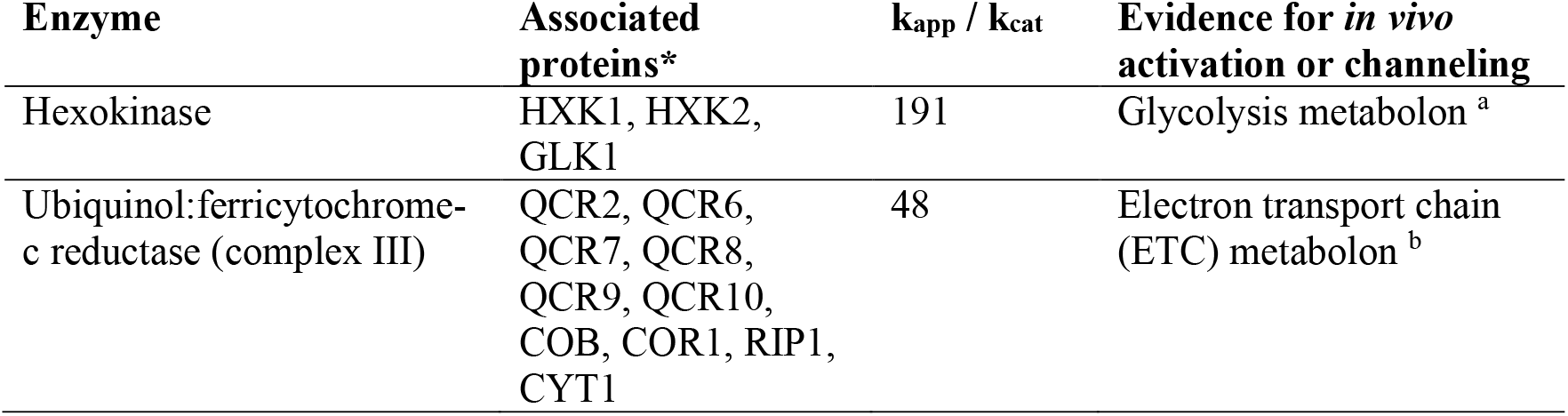
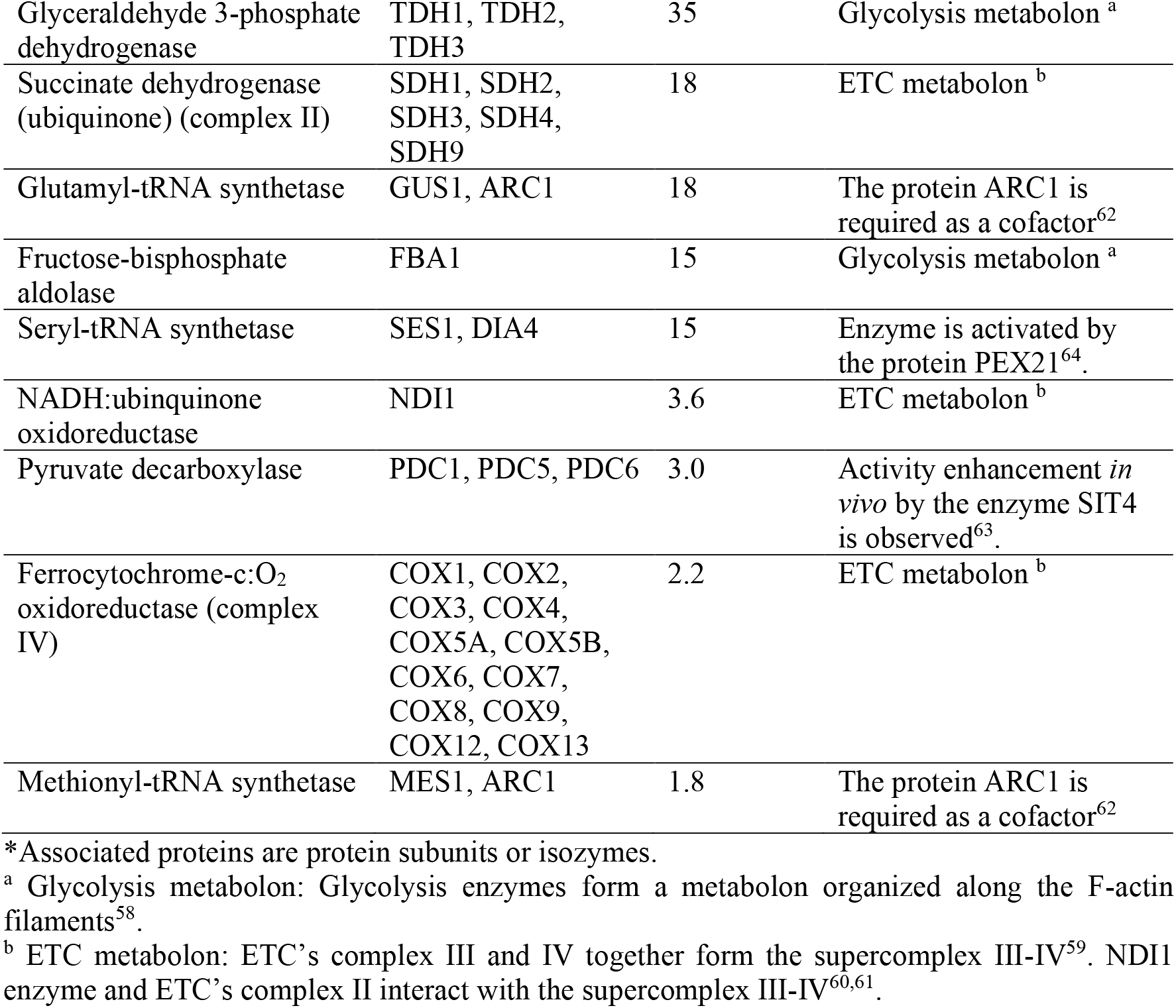
List of enzymes whose k_app_ exceed k_cat_ values with independent experimental confirmation.

| Enzyme | Associated proteins* | $k_{app} / k_{cat}$ | Evidence for *in vivo* activation or channeling |
| --- | --- | --- | --- |
| Hexokinase | HXK1, HXK2, GLK1 | 191 | Glycolysis metabolon <sup>a</sup> |
| Ubiquinol:ferricytochrome-c reductase (complex III) | QCR2, QCR6, QCR7, QCR8, QCR9, QCR10, COB, COR1, RIP1, CYT1 | 48 | Electron transport chain (ETC) metabolon <sup>b</sup> |
| Glyceraldehyde 3-phosphate dehydrogenase | TDH1, TDH2, TDH3 | 35 | Glycolysis metabolon <sup>a</sup> |
| Succinate dehydrogenase (ubiquinone) (complex II) | SDH1, SDH2, SDH3, SDH4, SDH9 | 18 | ETC metabolon <sup>b</sup> |
| Glutamyl-tRNA synthetase | GUS1, ARC1 | 18 | The protein ARC1 is required as a cofactor<sup>62</sup> |
| Fructose-bisphosphate aldolase | FBA1 | 15 | Glycolysis metabolon <sup>a</sup> |
| Seryl-tRNA synthetase | SES1, DIA4 | 15 | Enzyme is activated by the protein PEX21<sup>64</sup>. |
| NADH:ubiquinone oxidoreductase | NDI1 | 3.6 | ETC metabolon <sup>b</sup> |
| Pyruvate decarboxylase | PDC1, PDC5, PDC6 | 3.0 | Activity enhancement *in vivo* by the enzyme SIT4 is observed<sup>63</sup>. |
| Ferrocytochrome-c:O<sub>2</sub> oxidoreductase (complex IV) | COX1, COX2, COX3, COX4, COX5A, COX5B, COX6, COX7, COX8, COX9, COX12, COX13 | 2.2 | ETC metabolon <sup>b</sup> |
| Methionyl-tRNA synthetase | MES1, ARC1 | 1.8 | The protein ARC1 is required as a cofactor<sup>62</sup> |

\*Associated proteins are protein subunits or isozymes.
<sup>a</sup> Glycolysis metabolon: Glycolysis enzymes form a metabolon organized along the F-actin filaments<sup>58</sup>.
<sup>b</sup> ETC metabolon: ETC's complex III and IV together form the supercomplex III-IV<sup>59</sup>. NDI1 enzyme and ETC's complex II interact with the supercomplex III-IV<sup>60,61</sup>.

**Table 2.** List of enzymes whose k_app_ exceed k_cat_ values by at least 10-fold without experimental evidence for an enhancement mechanism.

| Enzyme | Associated proteins* | $k_{app} / k_{cat}$ |
| --- | --- | --- |
| Nucleoside diphosphate kinase | YNK1 | 130 |
| ATP synthase | ATP1, ATP2, ATP3, ATP4, ATP5, ATP6, ATP7, ATP8, ATP14, ATP15, ATP16, ATP17, ATP18, ATP19, ATP20, OLI1, TIM11 | 68 |
| Isoleucyl-tRNA synthetase | ILS1 | 52 |
| L-Alanine transaminase | ALT1 | 44 |
| Inorganic diphosphatase | PPA2 | 35 |
| Histidyl-tRNA synthetase | HTS1 | 27 |
| Homocitrate synthase | LYS21 | 27 |
| Pyruvate carboxylase | PYC1, PYC2 | 26 |
| Valyl-tRNA synthetase | VAS1 | 22 |
| Glycyl-tRNA synthetase | GRS1, GRS2 | 19 |
| Cystathionine gamma-lyase | CYS3 | 15 |

\*Associated proteins are protein subunits or isozymes.

**Fig 3.**
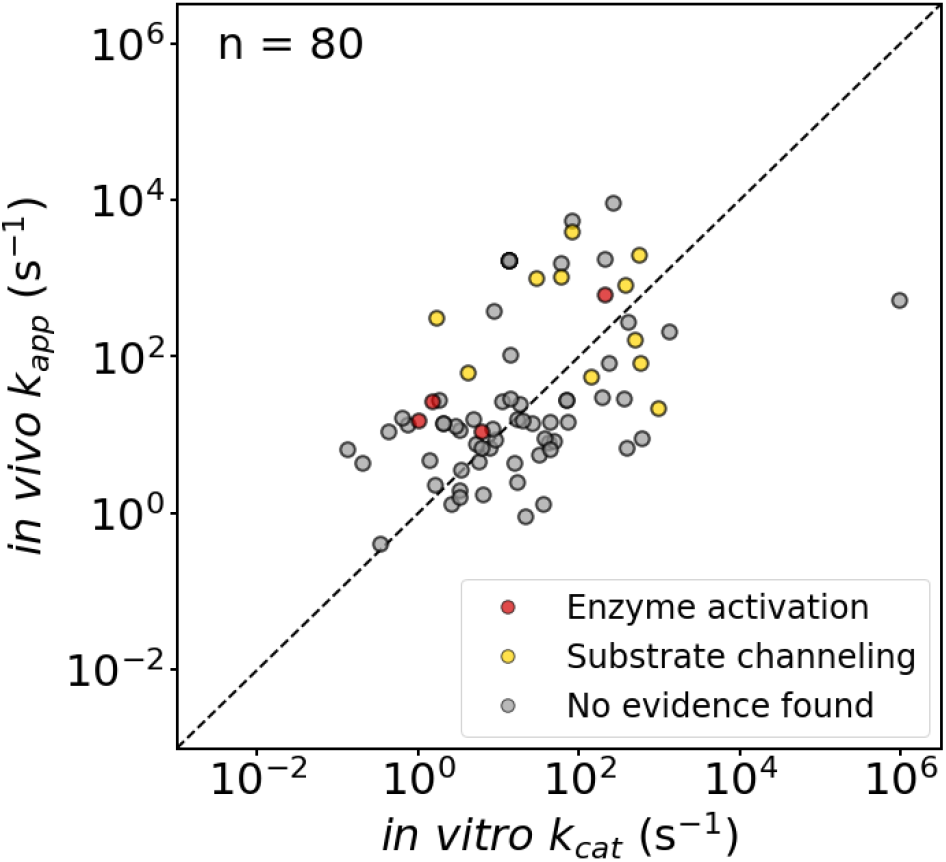
Comparison of *in vivo* k_app_ vs. *in vitro* k_cat_ values. k_app_ values for the growth condition of batch, aerobic, glucose, and minimal (YNB) media were used in comparison. Experimentally validated enzymes with enhanced *in vivo* efficiency are indicated by red and yellow dots. Gray dots indicate enzymes for which we did not identify such information.

k_app_ parameter sets were subsequently regressed separately under different growth conditions (see Methods) to assess their impact on enzyme efficiency. Perturbation instances from the reference condition (i.e., batch, aerobic, glucose, minimal (YNB) media) include (i) from aerobic to anaerobic, (ii) from minimal to rich (YNB + amino acids) media, (iii) from batch to chemostats (C- or N-limited), and (iv) from glucose to an alternative carbon substrate. Plots of 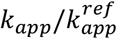 ratio values are shown in Fig. 4. 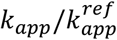 values reflect increased, constant, or reduced enzymatic efficiency under the perturbed from the reference condition. Overall, we found that replacing glucose with an alternative substrate leads to on an average as much as 85% reduction in enzymatic performance (i.e., when switching to trehalose) (see Fig. 4). This enzymatic performance reduction is mirrored in the observed slower growth rates^37^ (GR) on galactose (GR = 0.17 h^−1^), maltose (GR = 0.28 h^−1^), and trehalose (GR = 0.05 h^−1^) compared to (GR = 0.42 h^−1^) for glucose^30^. *sc*RBA results allude to likely strong adaptation of *S. cerevisiae* for proteome-efficient growth on glucose but not for the other three sugar substrates. Specifically, under galactose growth, galactose 1-phosphate accumulates while the fructose 6-phosphate pool is depleted^36^ suggesting the formation of a metabolic bottleneck between: (i) UDP-glucose – hexose-1P uridylyltransferase, (ii) UDP-glucose 4-epimerase, or (iii) phosphoglucomutase. Under maltose utilization, the maltose/proton symporter (uptake) and the alpha-glucosidase (maltose to glucose reaction) are likely rate-limiting as *S. cerevisiae* is susceptible to maltose hypersensitivity shock^119^. Under trehalose growth, the turnover number of the first enzymatic step, trehalase (k_app_ of 355 s^−1^), is comparable to the one for the glucose growth, hexokinase (k_app_ of 319 s^−1^). However, a closer look at the measured enzyme concentrations^30,37^ reveals an approximately 60-fold lower availability of trehalase (i.e., 0.22 nmol enzyme gDW^−1^) compared to hexokinase (i.e., 14 nmol enzyme gDW^−1^) for trehalose and glucose growth, respectively.

**Fig. 4.**
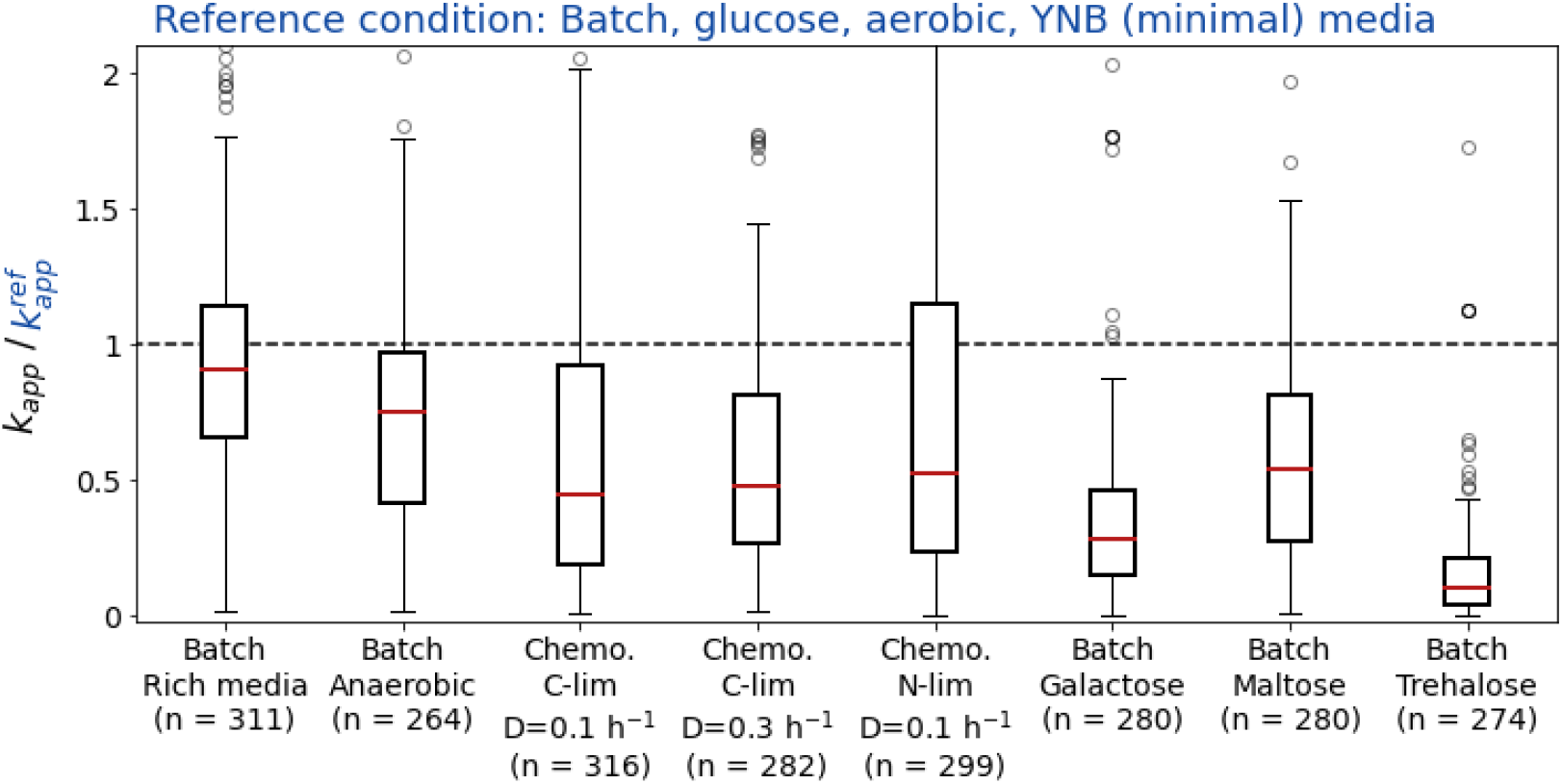
Distribution of estimated *in vivo* k_app_ ratios between perturbed and reference conditions. Distributions are visualized using standard box plots (i.e., box: interquartile range (IQR), whiskers: 1.5*IQR, and dots: outliers). The value of “n” indicates the number of overlapping reactions of the respective two sets of reactions whose k_app_ can be estimated from available flux and protein measurements.

Lack of oxygenation and nutrient limitation generally negatively affect enzymatic efficiency (see Fig. 4). Under C- and N-limitation, network-wide efficiency reduction is likely due to depleted intracellular metabolite pools^56^ which introduce substrate level limitations for many enzymes. Under anaerobic conditions, two-fold efficiency reduction (i.e., 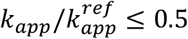) are predicted for enzymes in the TCA cycle performing biosynthesis role, ethanol fermentation (i.e., specifically alcohol dehydrogenase), fatty acid biosynthesis and elongations, and nucleotide biosynthesis. Amino acid supplementation to the (minimal) YNB media does not appear to significantly affect enzyme efficiency (see Fig. 4). Overall, using model *sc*RBA, we found that by estimating condition-specific *in vivo* k_app_ values we can elucidate changes in overall enzymatic efficiency utilization as a function of growth conditions and extracellular nutrient availability.

### *sc*RBA model simulation of Crabtree effect

We next contrasted *sc*RBA model predictions against experimental growth phenotypes at varying glucose uptake rates^29,30,33–37,39,41–44,47,120,121^. GAM_RBA_, NGAM, and *in vivo* k_app_ values for batch aerobic conditions were used in the simulations. *S. cerevisiae* expresses Crabtree-negative phenotype (i.e., no ethanol overflow) at low glucose uptake rates but switches to the Crabtree-positive phenotype (i.e., ethanol overflow) at high glucose uptake rates (see Fig. 5 for the data points from a variety of sources^29,30,33–37,39,41–44,47,120,121^). Comparisons of flux and proteomic data of respiration- and fermentation-favoring yeasts have implicated before ribosome availability and mitochondrial proteome bottlenecks as drivers of ethanol overflow. Here we performed RBA simulations with different combinations of capacity constraints to infer which limitation is actively limiting growth phenotypes at different levels of glucose uptake (see Fig. 5 for RBA simulation results). We denote with constraint (1) the overall proteome requirement as a % of the maximum, with constraint (2) the ribosome availability as a % of the total rRNA needed to complete all protein synthesis, and with constraint (3) the total mitochondrial protein capacity set to be less than 5% of the total proteome.

**Fig. 5.**
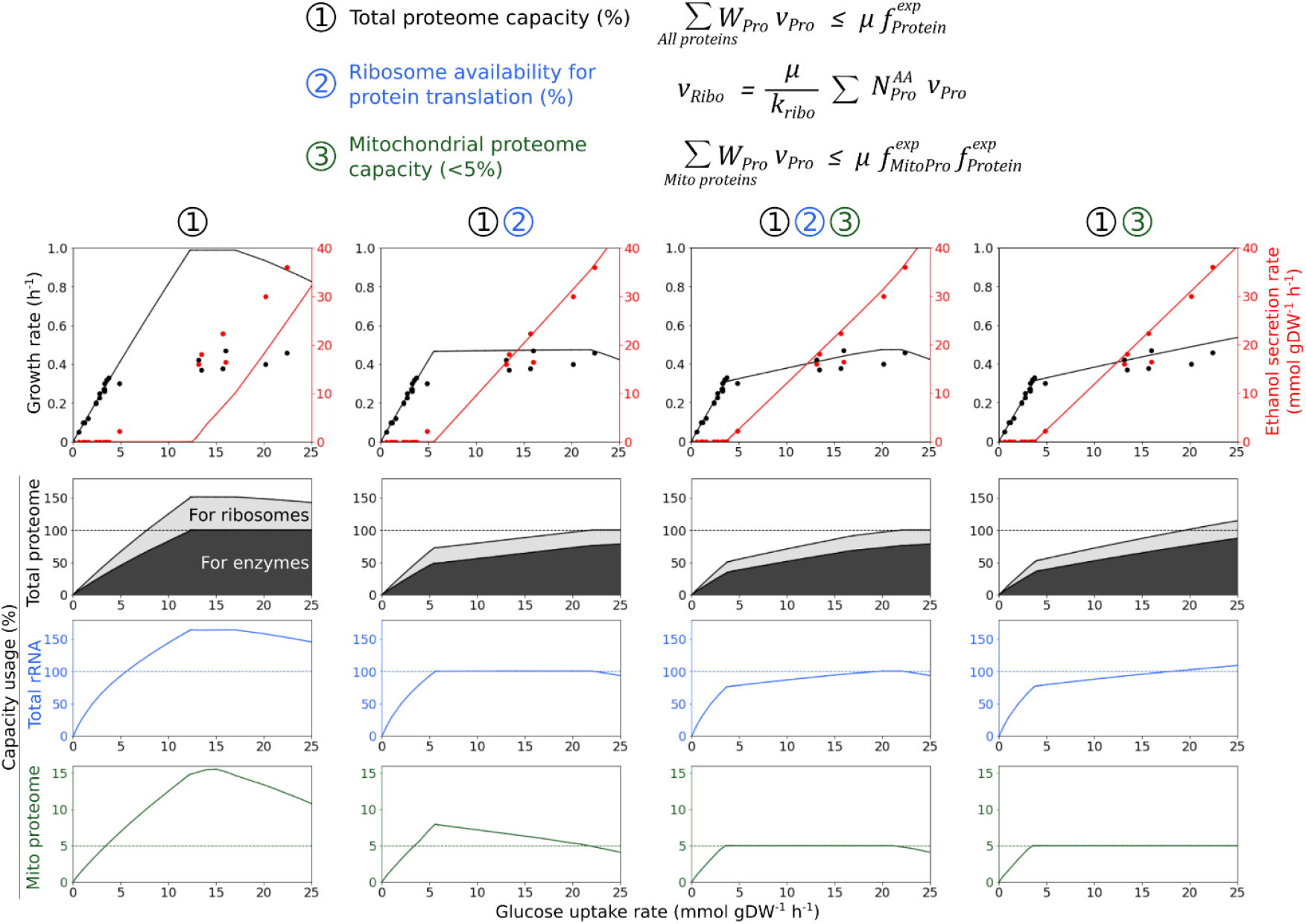
Model-simulated ethanol overflow phenotypes and capacity usage subject to proteome and ribosome limitations. Each column consists of figures reporting results with constraints being included in or excluded from RBA optimization as labeled. Figures in the top row report the simulated (as lines) and experimental (as dots) growth and ethanol secretion rates at different glucose uptake rates. Figures in the bottom three rows report predicted total proteome, total rRNA, and mitochondrial proteome capacity usage, respectively. In RBA runs lacking the ribosome availability constraint (2), ribosomal protein and rRNA capacity usage were back-calculated from the protein translation fluxes.

Starting with the first column of Fig. 5, RBA simulations lacking both ribosome and mitochondrial proteome availability constraints (2,3) predicted a significantly overestimated maximal growth rate (i.e., predicted 0.99 vs. observed 0.47 h^−1^) alongside a delayed overflow metabolism at much higher glucose uptake rates (i.e., 12.9 vs. at less than 4.8 mmol gDW^−1^ h^−1^ observed experimentally). Looking at the relevant plots (in the 1^st^ column of Fig. 5) we see that both rRNA and mitochondrial proteome capacity limitations are exceeded for glucose uptake rates greater than 5.6 mmol gDW^−1^ h^−1^ leading to predicted phenotypes that violate physiological capacity limits even for the total proteome (by up to 51%) due to the unaccounted cost of ribosomal proteins. In addition, the unaccounted in the model total required rRNA exceeds by up to 64% the experimental limit, whereas the unconstrained mitochondrial proteome reached up to 15.5% of the total proteome violating the 5% experimental limit. At very high glucose uptake rates (i.e., greater than 17 mmol gDW^−1^ h^−1^) the fraction of total proteome allocated to glucose transporter and catabolic enzymes ends up ultimately limiting overall growth. These results indicate that using only total proteome capacity limit fails to capture the true maximal growth rate and the onset of overflow metabolism.

The addition of the ribosome availability constraint (2) (i.e., see the 2^nd^ column of Fig. 5) resulted in RBA predictions that quantitatively match the experimental maximal growth rate as the cost of the protein fraction of the ribosomes is now correctly accounted for in the total proteome allotment. However, the onset of overflow metabolism is slightly delayed at a glucose uptake rate of 5.6 mmol gDW^−1^ h^−1^ compared to the experimental observation (i.e., at less than 4.8 mmol gDW^−1^ h^−1^). In addition, the growth rate reaches its maximum at an earlier glucose uptake rate of 5.6 mmol gDW^−1^ h^−1^ compared to the gradual increase seen in the experimental data points before the dip at very high glucose uptake rates due to the increased translation requirements for the extra glucose transporters and glycolysis. Note that this rapid plateauing of the growth rate is associated with a violation of the mitochondrial proteome capacity limit (i.e., 7.9% proteome usage vs. the 5% limit). Overall, the inclusion of the ribosome availability constraint (2) goes a long way towards recapitulating the physiological phenotype with only small departures.

Finally, adding the mitochondrial proteome capacity constraint (3) to the RBA model resulted in predictions that exactly matched (i) experimental maximal growth rates, (ii) the onset of overflow metabolism, and (iii) the more gradual increase of growth rate with glucose uptake (i.e., see 3^rd^ column of Fig. 2). Overflow metabolism was initiated at a glucose uptake rate of 3.7 mmol gDW^−1^ h^−1^ which is exactly the point at which the 5% mitochondrial proteome limit was reached. Ethanol overflow increased linearly with the glucose uptake rate which is consistent with growth-coupled metabolism required for NAD^+^/NADH rebalancing and ATP supply for growth. Total rRNA capacity was exhausted when the glucose uptake rate reached 20 mmol gDW^−1^ h^−1^ which matches exactly the point of maximum growth. Any further increase in glucose uptake rate was associated with a reduction in the maximum growth rate as the expanded demands for transporter and glycolysis enzymes claimed proteome capacity for more ribosomes to sustain faster growth.

To further delineate the impact of each constraint, we also reran the RBA model with the mitochondrial proteome but not the ribosome availability constraint (i.e., see the 4^th^ column of Fig. 5). The results are nearly identical with the ones with all three constraints present (i.e., 3^rd^ column in Fig. 5) until a glucose uptake of 20 mmol gDW^−1^ h^−1^ where the decrease in the growth rate is not observed. This is because at that point ribosome capacity and not mitochondrial proteome becomes the limiting factor in constraining growth due to competition against ribosomes for proteome capacity from glucose transporter and glycolysis. These results imply that the effect of ribosome availability is subsumed within the mitochondrial capacity constraint till a high glucose uptake rate of 20 mmol gDW^−1^ h^−1^ is reached requiring the direct incorporation of the ribosomal availability constraint.

Overall, simulation results from model *sc*RBA suggested that mitochondrial proteome capacity is exhausted before rRNA capacity triggering ethanol overflow. Nevertheless, it appears that both rRNA and total proteome capacity usage percentages at the maximum growth rate (i.e., 100% and 97%, respectively) suggest that rRNA molecules are produced in (near) proportion to physiological quantities of mitochondrial and total proteome in *S. cerevisiae* so as not to under-utilize any macromolecules which are costly to synthesize. This raises the question of why mitochondrial proteome and rRNA capacity in *S. cerevisiae* are not expanded to facilitate even faster growth rates. For example, *Escherichia coli* has a much larger RNA capacity (i.e., 20.5% ^122^ vs. 6.6% gRNA gDW^−1^ in *S. cerevisiae*^74^) and a slightly more efficient ribosome (i.e., k_ribo_ of 17 ^123^ vs. 13.2 in *S. cerevisiae* (this work)). This enables *E. coli* to access a significantly higher maximal growth rate than *S. cerevisiae* (i.e., max growth of 1.2 h^−1 124^ vs. 0.49 h^−1^ in *S. cerevisiae*^30^). Correlation between low abundance of mitochondrial proteome fractions and ethanol overflow phenotypes has been observed in proteomics and flux data^111,125^ of several Crabtree-positive and negative yeast species. Correspondingly, Crabtree-negative yeasts grow faster than *S. cerevisiae* and other Crabtree-positive yeasts under aerobic conditions^111,125,126^. While at first glance this is counter-intuitive, the slower growing ethanol overflow phenotype offers a number of evolutionary advantages: (i) out-competing other microbes in sugar consumption, (ii) producing ethanol that is toxic to bacterial competitors, and (iii) adapting easily to anaerobic conditions with ready to deploy respiro-fermentative proteome allocation^111,127^. The adaptive advantage of *S. cerevisiae*’s proteome was demonstrated in anaerobic and cyclically oxygenated cultures where higher abundances of *S. cerevisiae* cells competing with the respiratory-favoring yeast *Issatchenkia orientalis* were measured^111^. Overall, model *sc*RBA results corroborate literature-reported observations and strongly suggest that mitochondrial proteome limitation first triggers overflow metabolism whereas rRNA capacity ultimately lowers maximum growth rate at very high uptake rates.

### Effect of protein capacity limit on metabolic flux upper bounds and maximal product yields

Enzyme(s) availability bottlenecks can add additional barriers to reaching FBA calculated maximum theoretical limits. Identifying these yield-limiting enzymes is important so as to guide specific gene overexpression strategies remedying these shortcomings without wasting resources on enzymes that are not limiting. To this end, we contrasted the calculated flux bounds (i.e., FVA analysis) using model *sc*RBA (with k_app_ parameters for batch aerobic conditions typically used in compound production) and model *iSace*1144 using FBA. RBA/FBA absolute upper bound flux ratios were calculated for 800 flux-carrying metabolic reactions under glucose uptake conditions. We performed RBA runs with and without the mitochondrial proteome capacity constraint (i.e., limiting to 5% of total proteome capacity) and contrasted results to pinpoint mitochondrially originated metabolic limitations (see Fig. 6 for a summary and see Supplementary Data 5 for details).

**Fig. 6.**
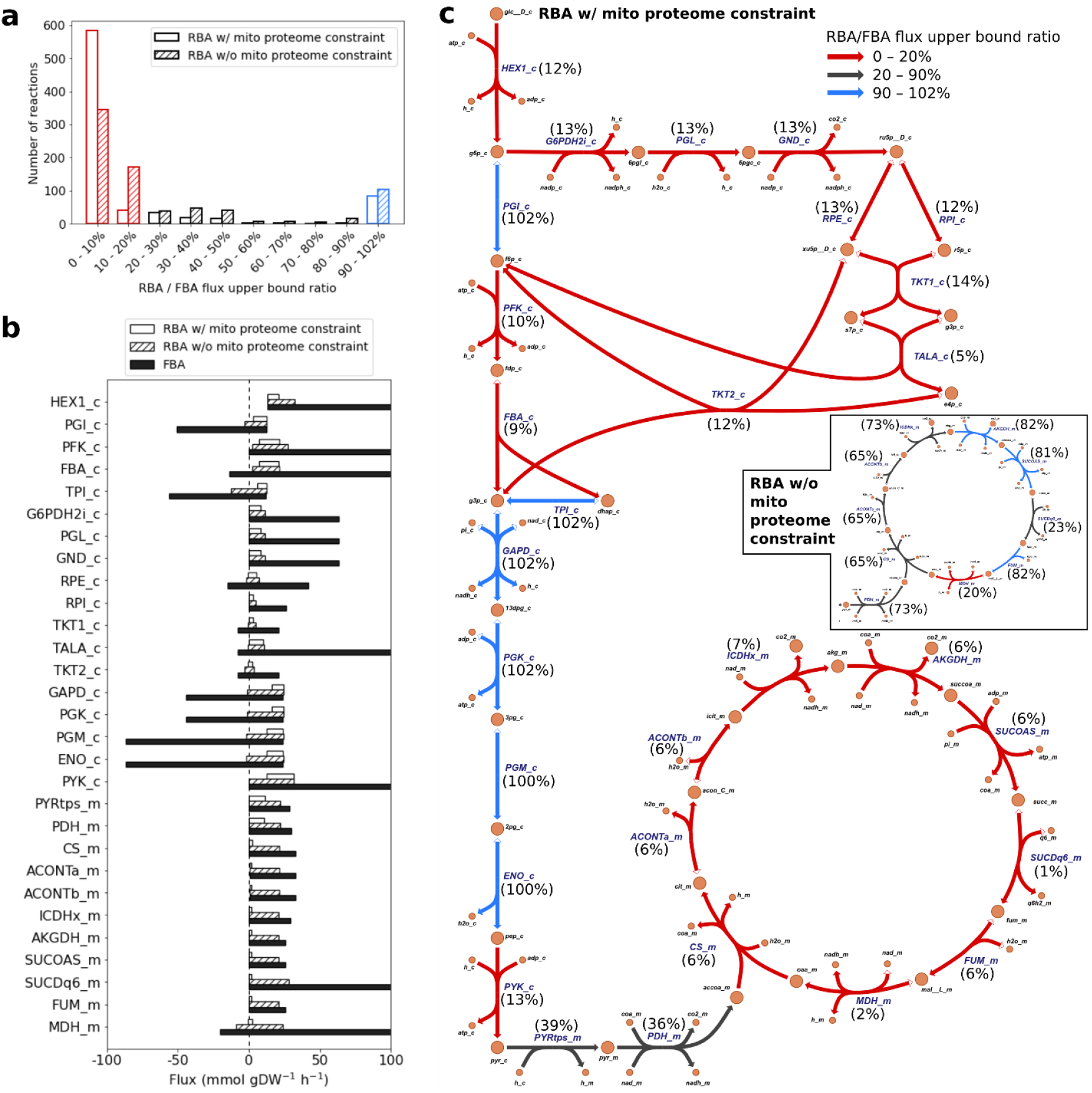
*sc*RBA model’s metabolic flux variability analysis accounting for protein capacity limit. (**a**) Histogram of RBA/FBA flux upper bound ratio values. (**b**) RBA- (white and striped bars) and FBA-calculated (black bars) flux ranges for reactions in central metabolism subject to experimental glucose uptake and growth rates^30^. Reaction IDs are in BiGG format^128^ and reaction details are available in the *sc*RBA github repository. (**c**) Central metabolism network (drawn by the Escher software^129^) with overlayed reaction IDs and corresponding RBA/FBA flux upper bound ratio values (annotated as colors of arrows) for RBA runs with mitochondrial proteome constraint. The figure inset shows much higher flux upper bounds for TCA cycle reactions from RBA runs without the mitochondrial proteome constraint.

Under only total proteome and rRNA capacity limitations, RBA/FBA ratios are less than 20% for as many as 516 out of 800 flux-carrying reactions (see Fig. 6a). This indicates that catalytic resource limitation as encoding in model *sc*RBA are propagated to most reactions in the metabolic network. In central metabolism, FBA (through FVA analysis) allows for maximal glycolysis and pentose phosphate pathway (PPP) fluxes that are up to an order of magnitude larger than the glucose uptake rate (i.e., 13.2 mmol gDW^−1^ h^−1^) (see Fig. 6b). These very high fluxes are caused by activating ATP-consuming cycles. For example, the FBA-calculated maximal flux of phosphofructosekinase (ID: PFK_c) reaction in glycolysis is 225 mmol gDW^−1^ h^−1^ which contains an ATP-consuming cycle (i.e., 95% of total flux) with fructose bisphosphate phosphatase reaction in gluconeogenesis. These cycles are retained in FBA because without any additional constraints extra glucose can be used to produce ATP at a yield of up to 25.6 mol ATP / mol glucose. Imposing the total proteome capacity constraint in RBA greatly reduce the extent of ATP-consuming cycles (see Fig. 6b and 6c). For example, the RBA-calculated PFK_c maximal flux is 27.6 mmol gDW^−1^ h^−1^ which is an order of magnitude smaller than the FBA-calculated one.

Adding the mitochondrial proteome capacity constraint significantly reduces the RBA/FBA flux upper bound ratios of the TCA cycle reactions from 80% to 6% (see Fig. 6c) and of electron transport chain (ETC) from 99% to 24%. FBA-predicted TCA and ETC fluxes are much higher than RBA-predicted values because of the readily available precursors acetyl-CoA and NADH (respectively) synthesized from the glucose uptake surplus. Reduced mitochondrial protein capacity for respiration leads to lowered ATP availability and ultimately constrains ATP-consuming flux cycles reflected in the significantly lower RBA/FBA flux upper bound ratios for many reactions across the metabolic network (see Fig. 6a). Overall, the RBA framework implementing total proteome capacity constraint provides flux predictions with large reductions for fluxes that can participate in futile ATP-consuming cycles. Implementing mitochondrial proteome capacity constraints has a direct and dramatic impact on the maximal fluxes of mitochondrial enzymes and an indirect effect on non-mitochondrial enzymes constrained by ATP availability.

Fluxes of the six non-cycling glycolysis reactions are well resolved through FBA as they are fully coupled to the pre-specified glucose uptake (see Fig. 6c). Counterintuitively, their upper bounds are slightly higher using RBA (with or without the mitochondrial proteome capacity constraint) than using FBA by about 1.5 – 1.8% (see Fig. 6c). This is because a slightly higher glycolysis/pentose phosphate pathway (PPP) split ratio for a given glucose uptake is predicted by optimizing NADPH usage in the *sc*RBA model. The lower flux through the NADPH-generating PPP is due to the fact that the actual NADPH needs for amino acid synthesis (accounting in detail by RBA) is slightly less than the lumped amount of the stoichiometric description used in FBA.

Next, we evaluated the effect of protein capacity limits on the production yields for 28 compounds by contrasting their maximal yields from *sc*RBA and *iSace*1144 models (see Table 3 and Supplementary Data 5 for RBA-predicted fluxes and proteome allocations). Separate RBA simulations with and without the mitochondrial proteome capacity constraint (i.e., < 5% of total proteome) were performed to identify metabolites whose yields are negatively affected under mitochondrial proteome limitations.

**Table 3.** FBA-predicted, RBA-predicted, and literature-reported experimental yields for 28 products.

| Product | Precursors | $Y_{p,max}^{FBA}$ | $Y_{p,max}^{RBA}^{**}$ | | $Y_{p,max}^{RBA} / Y_{p,max}^{FBA}^{**}$ | | $Y_p^{exp\#\#}$ | Pathway & yield reference |
| --- | --- | --- | --- | --- | --- | --- | --- | --- |
|  |  |  | w/o MP | w/ MP | w/o MP | w/ MP |  |  |
| Free fatty acid | Fatty acyl-CoA | 0.30 | 0.26 | 0.26 | 87% | 87% | 0.1 | 83 |
| Fatty alcohol | Fatty acyl-CoA | 0.27 | 0.24 | 0.24 | 88% | 88% | 0.005 | 7 |
| Triacylglycerol<sup>++</sup> | Triacylglycerol | 0.29 | 0.26 | 0.26 | 88% | 88% | 0.14 | 94 |
| Citramalate | Acetyl-CoA and pyruvate | 0.77 | 0.77 | 0.77 | 100% | 100% | N/A | 103 |
| n-Butanol | (Acetyl-CoA and pyruvate) or L-threonine | 0.35 | 0.32 | 0.32 | 91% | 91% | 0.04 | 103 |
| Polyhydroxybutyrate | Acetoacetyl-CoA | 0.54 | 0.52 | 0.52 | 97% | 97% | 0.13 | 104 |
| Artemisinic acid | Farnesyl diphosphate | 0.28 | 0.23 | 0.23 | 83% | 83% | N/A | 105 |
| Sesquiterpene | Farnesyl diphosphate | 0.31 | 0.26 | 0.26 | 84% | 84% | 0.003 | 106,107 |
| Ethanol<sup>++</sup> | Ethanol | 0.49 | 0.48 | 0.48 | 100% | 100% | 0.44 | 108 |
| L-Lactate | Pyruvate | 0.91 | 0.90 | 0.90 | 100% | 100% | 0.69 | 109 |
| (2R,3R)-Butanediol | Pyruvate | 0.49 | 0.49 | 0.49 | 100% | 100% | 0.11 | 84 |
| **Isobutanol** | **Isobutanol** | **0.39** | **0.34** | **0.05** | **86%** | **12%** | **0.06** | 85 |
| 3-Hydroxypropionic acid | Malonyl-CoA | 0.76 | 0.69 | 0.69 | 91% | 91% | 0.13 | 86 |
| **3-Hydroxypropionic acid** | **L-Alanine** | **0.84** | **0.81** | **0.49** | **96%** | **58%** | **0.14** | 87 |
| Succinate<sup>++</sup> | Succinate | 0.93 | 0.20 | 0.20 | 21% | 21% | 0.11 | 88,130 |
| **Malate<sup>++</sup>** | **Malate** | **1.18** | **1.16** | **1.07** | **98%** | **91%** | **0.31** | 89 |
| Shikimate<sup>++</sup> | Shikimate | 0.71 | 0.70 | 0.70 | 99% | 99% | N/A | 90 |
| cis,cis-Muconate | 3-Dehydroshikimate | 0.61 | 0.60 | 0.60 | 99% | 99% | 0.07 | 91 |
| **p-Hydroxybenzoate** | **Chorismate** | **0.52** | **0.51** | **0.50** | **98%** | **95%** | **0.07** | 92 |
| **p-Aminobenzoate<sup>++</sup>** | **p-Aminobenzoate** | **0.50** | **0.42** | **0.41** | **85%** | **82%** | **0.01** | 93 |
| 2-Phenylethanol | L-Phenylalanine | 0.36 | 0.35 | 0.35 | 99% | 99% | 0.08 | 95 |
| Styrene | L-Phenylalanine | 0.30 | 0.29 | 0.29 | 98% | 98% | 0.001 | 96 |
| p-Coumaric acid | L-Tyrosine | 0.49 | 0.48 | 0.48 | 98% | 98% | 0.15 | 97 |
| Reticuline | L-Tyrosine | 0.35 | 0.24 | 0.24 | 68% | 68% | N/A | 98 |
| (2S)-Naringenin | L-Tyrosine and malonyl-CoA | 0.45 | 0.43 | 0.43 | 96% | 96% | N/A | 99 |
| Resveratrol | L-Tyrosine and malonyl-CoA | 0.38 | 0.36 | 0.36 | 96% | 96% | 0.0002 | 100 |
| Glycerol<sup>++</sup> | Glycerol | 0.77 | 0.75 | 0.75 | 97% | 97% | 0.5 | 101 |
| **1,3-Propanediol** | **Glycerol** | **0.57** | **0.49** | **0.47** | **86%** | **82%** | **N/A** | 102 |

Optimization constraints: growth rate = 0.1 h<sup>-1</sup>, glucose uptake rate's upper bound = 13.2 mmol gDW<sup>-1</sup> h<sup>-1</sup> (referencing experimental value<sup>30</sup>). $Y_{p,max}^{FBA}$ and $Y_{p,max}^{RBA}$ (in g g-Glucose<sup>-1</sup>): FBA- and RBA-calculated maximal theoretical yields. $Y_p^{exp}$ : literature-reported experimental yields.
<sup>\*\*</sup>: Separate RBA yield values were calculated with mitochondrial proteome capacity constraint included and excluded (i.e., labeled with “w/ MP” and “w/o MP”, respectively). Rows are in bold-face if calculated yields with mitochondrial proteome constraint are different than the ones without.
<sup>++</sup>: These metabolites are produced with native pathways
<sup>##</sup>: “N/A”, or not available, is designated to metabolites with experimental studies where only titers were reported. Citramalate and shikimate were intermediates for other terminal metabolite productions and thus their yields were not reported. A higher yield of 0.43 g-succinate g-glucose<sup>-1</sup> was reported for a laboratory-evolved strain.

Under only total proteome and rRNA capacity limitations, the maximal yields for 26 out of 28 product metabolites are only marginally restricted with RBA-calculated yields well within 80-100% of the FBA-calculated values (see Table 3). Butanediol, citramalate, ethanol, and lactate productions are least affected with RBA-predicted yields retained at 100% of FBA-calculated yields. This is consistent with the previously determined high protein reserve capacity of *S. cerevisiae* for overflow metabolism. In contrast, succinate and reticuline maximal yields are significantly affected by the total proteome capacity limit (i.e., down to 21% and 68% of FBA-calculated yields, respectively). Succinate production is directly limited by the enzyme efficiency of its cytosolic pathway. We calculated for *S. cerevisiae* k_app_ values of 13, 30, 71, 1,582 s^−1^ for pyruvate-to-succinate enzymes compared to k_app_ values of 539 and 612 s^−1^ for pyruvate-to-ethanol enzymes. This kinetic bottleneck is absent in *E. coli* whose pyruvate-to-succinate enzyme k_cat_ values (archived in BRENDA^57^) are 250, 540, 931, and 1,150 s^−1^. Model *sc*RBA-predicted succinate yield (i.e., 0.20 g g-Glucose^−1^) matches the experimentally observed yield ranges for engineered *S. cerevisiae* strains ranging from 0.11 for an unevolved strain^88^ and 0.43 for a laboratory evolved strain^130^. These achieved succinate yields are much lower than the FBA-calculated maximum theoretical yield of 0.95 or alternatively that of an engineered *E. coli* strain (i.e., 0.81)^131^. As demonstrated in this succinate case study RBA results can provide a roadmap for relieving production bottlenecks through gene overexpression and/or replacement of enzymes with more efficient heterologous versions. The production of reticuline, on the other hand, is not limited by the efficiency of the enzymes in the biosynthetic pathway but rather by the proteome cost of recharging the needed cofactors (i.e., two NADPH and three S-adenosyl methionine (SAM) per reticuline molecule)^98^. Using model *sc*RBA we predicted that supplying NADPH (i.e., through PPP) for reticuline consumes only 1.6% of total available proteome. However, the recovery of SAM (C_1_ donor) from S-adenosyl homocysteine (SAH) for reticuline production is approximately 20-fold more proteome consuming than recovering NADPH taking up to 37% of the total proteome. The significant metabolic burden of recovering cofactor SAM has also been reported for *E. coli* where a 12-fold increase in reticuline titer was achieved when the dopamine-to-reticuline conversion occurred *ex vivo* with the growth medium was supplemented in excess with SAM^132^. The costliest product metabolite in terms of NADPH consumption is fatty alcohol requiring 16 moles of NADPH per mole of C_16_. Despite this high NADPH cost, the corresponding protein allocation for NADPH recharging is only 6.4% of the total proteome. *sc*RBA results demonstrate that a high demand for expensive-to-recharge cofactors can create significant proteome allocation burdens that can negatively affect maximal production yields. This suggests that more enzyme efficient cofactor recharging variants may be a promising route for de-bottlenecking flux through the product biosynthetic pathway.

Adding the mitochondrial proteome capacity constraint has little to no effect on the maximal yields for all products except isobutanol and 3-hydroxypropionic acid via the beta-alanine pathway consuming L-alanine as precursor (i.e., down to 12% and 58% of FBA-predicted yields, respectively) (see Table 3). This reduction is due to mitochondrial capacity limitations rather than ATP requirements. For both pathways a mitochondrial pyruvate transporter and various mitochondrial biosynthetic enzymes (i.e., three enzymes in isobutanol pathway and alanine transaminase in 3-hydroxypropionic acid pathway) are needed. ATP availability (mainly provided by mitochondrial respiration) is not the limiting factor for isobutanol and 3-hydroxypropionic acid production as no net ATP is needed starting from pyruvate (see Table 3). This is in contrast to biomass synthesis that has a much higher ATP demand and it is thus indirectly limited by mitochondrial proteome availability needed for ATP synthesis (see previously the Crabtree effect simulation results). In agreement with these remarks, relocating the mitochondrial enzymatic steps to cytosol and knocking out competitive pathways in an isobutanol overproducing *S. cerevisiae* strain resulted in a 200-fold yield improvement compared to wild-type^85^. Overall, RBA results demonstrate that limitations associated with compartment space (i.e., mitochondria herein) can negatively affect metabolite production with biosynthesis routes requiring (some) enzymes within the compartments. In contrast, ATP availability does not appear to be limiting under aerobic conditions.

Although protein capacity is not flagged as an issue for most metabolic products, experimentally achieved yields are usually significantly lower than RBA model predicted values except for ethanol and L-lactate (see Table 3). This underperformance may suggest that there is likely untapped potential to further optimize yield through the application of metabolic engineering strategies. Model *sc*RBA predicted proteome allocations for the maximal production of 28 metabolic products are provided in the Supplementary Data 6. These values can help guide further engineering strategies by contrasting with experimentally elucidated quantitative proteomic levels to better guide further strain redesign efforts.

## Discussion

Model *sc*RBA provides insights onto *S. cerevisiae* metabolism under a variety of growth conditions by imposing enzyme and ribosome availability limitations on top of mass balances and biomass synthesis needs. *sc*RBA was designed to reduce the number of variables and parameters to only the ones that can be supported by the available proteomic and fluxomic data. Biological processes not contributing to the functional parts of the model (e.g., transcriptional machinery) are modeled in an aggregate manner as sinks for redox and carbon resources without tracking individual steps (see Methods). This makes *sc*RBA more computationally tractable in analyzing metabolic proteome allocation compared to models of larger scope^27,28^. If details on transcription and mRNA availability are available then the ETFL framework^27^ provides an alternative. Whole-cell modeling^28^ has also been performed for *S. cerevisiae* but computational efficiency and missing parameter values (as noted by the authors) remain a challenge. A distinguishing feature of *sc*RBA is that parameters GAM, NGAM, and k_app_ are derived directly from *in vivo* data thus enabling the condition-specific prediction of growth yield, rate, and proteome allocation. We found significant variation in these parameters depending on nutrient and oxygen availability as well as carbon substrate choice driving the message that their values are condition-dependent. It is possible that parameter values may ultimately converge to the ones for the reference condition upon adaptation through laboratory evolution for different growth conditions, substrates, and/or genetic backgrounds as seen for *E. coli*^116^.

*sc*RBA results pinpoint that limitations in mitochondrial proteome and in rRNA capacity (for ribosome) are the primary and secondary barriers, respectively preventing yeast from growing faster under aerobic glucose uptake conditions. Reserve non-mitochondrial proteome capacity allows for ethanol overflow to be carried out even in the presence of oxygen. *In vivo* observations reveal that in *S. cerevisiae* the mitochondria per cell volume/total volume ratio can be as much as 35% under ethanol uptake conditions^78^. However, it remains limited to 5% under glucose uptake conditions^78^ suggesting that limiting mitochondrial proteome is not a maladaptation but rather a coordinated response activating the ethanol overflow program in *S. cerevisiae*. The predicted link between limited growth capacity and metabolic overflow in *S. cerevisiae* raises a question on its generality. Analogous overflow phenotypes are present for *E. coli* with acetate^124^ and cancer cells with lactate^133^ overflow. A quantitative framework similar to *sc*RBA could in principle be applied to confirm or refute the preserve of excess proteome at the point of respiratory or rRNA capacity exhaustion. It is important to stress that in addition to rRNA and proteome capacity bottlenecks examined in our RBA analysis transcriptional bottlenecks may be at play. Accounting for them would require *in vivo* data on RNA polymerase efficiency and mRNA availability.

The key design feature of *sc*RBA of using k_app_ as opposed to k_cat_ values unlocks the opportunity to parameterize many more reactions by leveraging the relative availability of quantitative proteomic and fluxomic datasets^29,30,34,37^. *sc*RBA makes use of 336 k_app_ values pegged to *in vivo* protein and flux measurements whereas only 80 k_cat_ values for *S. cerevisiae* under batch aerobic conditions with glucose as substrate can be recovered from BRENDA^57^. Using k_app_ values also obviates the need to “approximate” missing k_cat_ values with measurements from other organisms^134^. Contrasting *in vivo* k_app_ values with *in vitro* k_cat_ values can be an effective method for leveraging multi-omics data to systematically identify enzymes with *in vivo* enhancements. While the mechanism of enhancements may remain elusive, their presence and extent could reveal important biological insight as to the spatial organization of pathways and regulation of enzymes. Consistent with metabolomics data^36,56^ we observe a lowering in k_app_ values under substrate or nitrogen limitations caused by reduced metabolite concentrations. This further reinforces the need to re-estimate k_app_ parameters whenever simulation conditions change. An alternative approach could be to track enzyme turnover and enzyme saturation as carried out in the growth balance analysis (GBA) framework^135^. GBA uses formal (nonlinear) kinetic descriptions (e.g., Michaelis-Menten) for the reaction fluxes and mass balance equations for all macromolecules as in RBA. This enhanced level of detail in description comes at the expense of having to generate absolute intracellular metabolite concentration measurements (on top of flux and enzyme measurements) as inputs to estimate enzyme saturation parameters (e.g., K_M_ in the Michaelis-Menten equation). Therefore, even though RBA sacrifices the ability to directly model enzyme saturation, it can quantify proteome allocation at a genome-scale in a computationally and data efficient manner.

Model *sc*RBA is shown to be more apt at recapitulating *S. cerevisiae* metabolic phenotypes compared to the corresponding FBA model *iSace*1144^111^. In particular, *sc*RBA can both identify reaction step bottlenecks and often pinpoint the functional reason for them such as limited compartmental space, inefficient enzyme with low k_app_, or proteome costly cofactor cycling. We found that a significant fraction of *S. cerevisiae* reactions has an upper bound that is set correctly at a much lower value by *sc*RBA due to the mitochondrial and total proteome allocation limits. Despite this *sc*RBA-predicted product yield calculations are generally only marginally lower than the FBA-based theoretical limit. This is primarily due to the fact that most predicted upper bound excursions by FBA are associated with reactions that can participate in ATP-consuming cycles. While this ATP overhead is tolerated in FBA calculations by draining from a large glucose surplus, the associated increased proteome allotments are severely curtailed in RBA calculations. Because by design product synthesis pathways do not involve futile cycles, reductions in the predicted maximum yields under RBA are not significant. It is important to stress that the RBA modeling framework does not account for many other factors that could affect production such as regulatory feedback and metabolite pool bottlenecks^136,137^. For example, release of inhibition of several upstream enzymatic steps by high concentrations of tyrosine and phenylalanine need to be addressed when producing shikimate-derived metabolites in *S. cerevisiae*^90^. Overall, we envision *sc*RBA model as part of a combined and inter-operable experimental and computational toolkit to investigate cellular metabolism and assess metabolic proteome allocation of *S. cerevisiae*. The relative compact nature of the RBA framework presented herein makes it tractable for non-model organisms leveraging advances in GSM reconstructions^138^.

## Acknowledgements

We thank Patrick F. Suthers (from Penn State) for his comments. Computations for this research were performed on the Pennsylvania State University’s Institute for Computational and Data Sciences’ Roar supercomputer. This work was funded by the DOE Center for Advanced Bioenergy and Bioproducts Innovation (U.S. Department of Energy, Office of Science, Office of Biological and Environmental Research under Award Number DE-SC0018420). Any opinions, findings, and conclusions or recommendations expressed in this publication are those of the author(s) and do not necessarily reflect the views of the U.S. Department of Energy.

## Data availability

Data is available in the GitHub repository https://github.com/maranasgroup/scRBA.

## Code availability

Software scripts are available in the GitHub repository https://github.com/maranasgroup/scRBA.

The user manual is provided in the Supplementary Methods.

## Author contributions

Conceptualization: H.V.D. and C.D.M; methodology, software, and data curation: H.V.D.; analysis: H.V.D. and C.D.M.; resources, supervision, and funding acquisition: C.D.M.; writing: H.V.D. and C.D.M.

