## Supplementary Methods for "Evaluating proteome allocation of *Saccharomyces cerevisiae* phenotypes with resource balance analysis"

### A. Mathematical formulation

In this section, a formulation overview is first provided. Then, we provide all necessary details for one to reconstruct the yeast RBA model and associated constraints. We will start with “Section A.1. Sets, indexes, and variables” which establish the starting mathematical structure necessary for equations and constraints in Section A.2 – A.4. The equations and constraints in the model are boxed whereas the reaction and mathematical equations used in derivation and explanation are not. Model stoichiometric coefficients and parameters are mentioned alongside with the equations and constraints that they are in (throughout Section A.2 – A.4).

The model *sc*RBA consists of (macro)molecules and reactions for metabolism and cellular machinery production linked through steady-state mass balance constraints as in FBA ^1^ (see Figure S1.1 for a schematic representation).

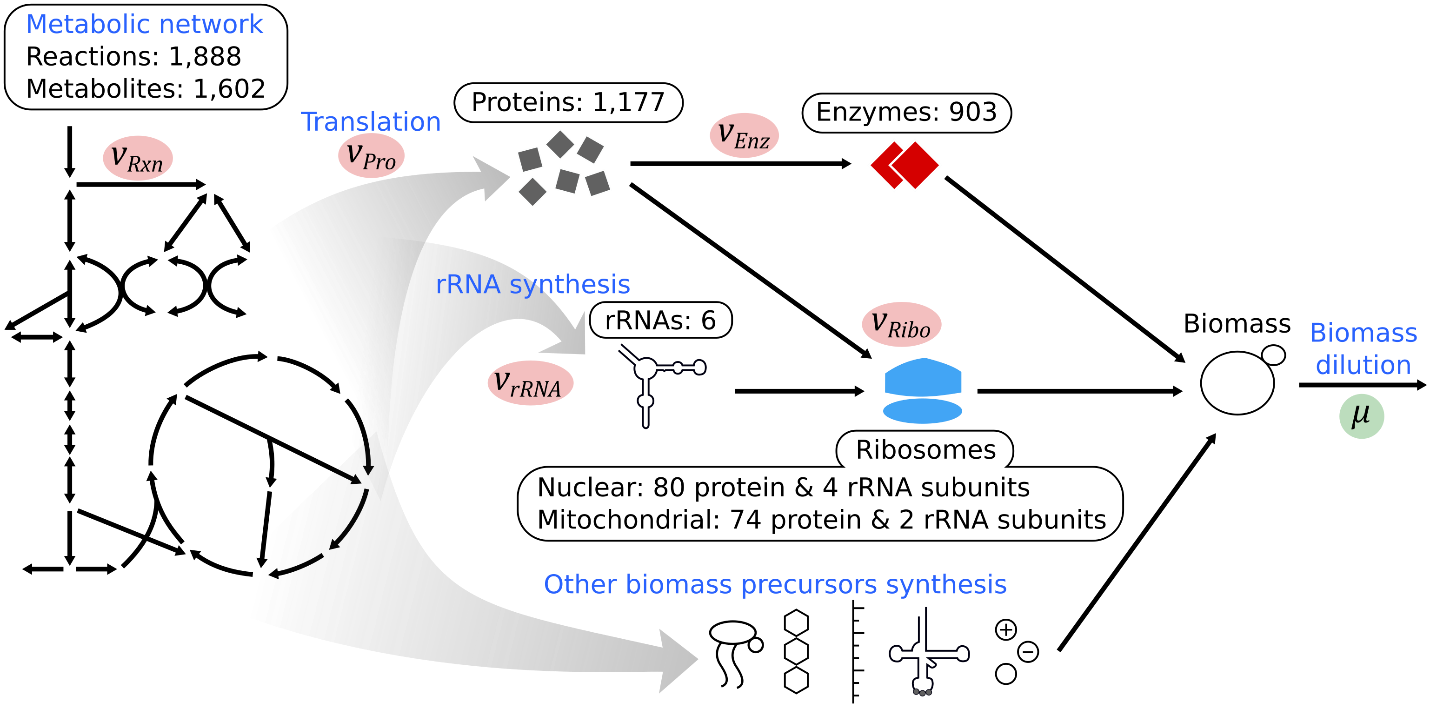

**Figure S1.1**. Overview of *sc*RBA mass balance.

An overview of the RBA optimization model that identifies the maximal growth rate is provided in Figure S1.2. By fixing the growth rate the *sc*RBA model is converted into a linear programming LP formulation (i.e., RBA-LP) which can be efficiently solved. A bisection method is employed to iteratively collapse the estimated (infeasible) upper and (feasible) lower growth rates until they are within the tolerance.

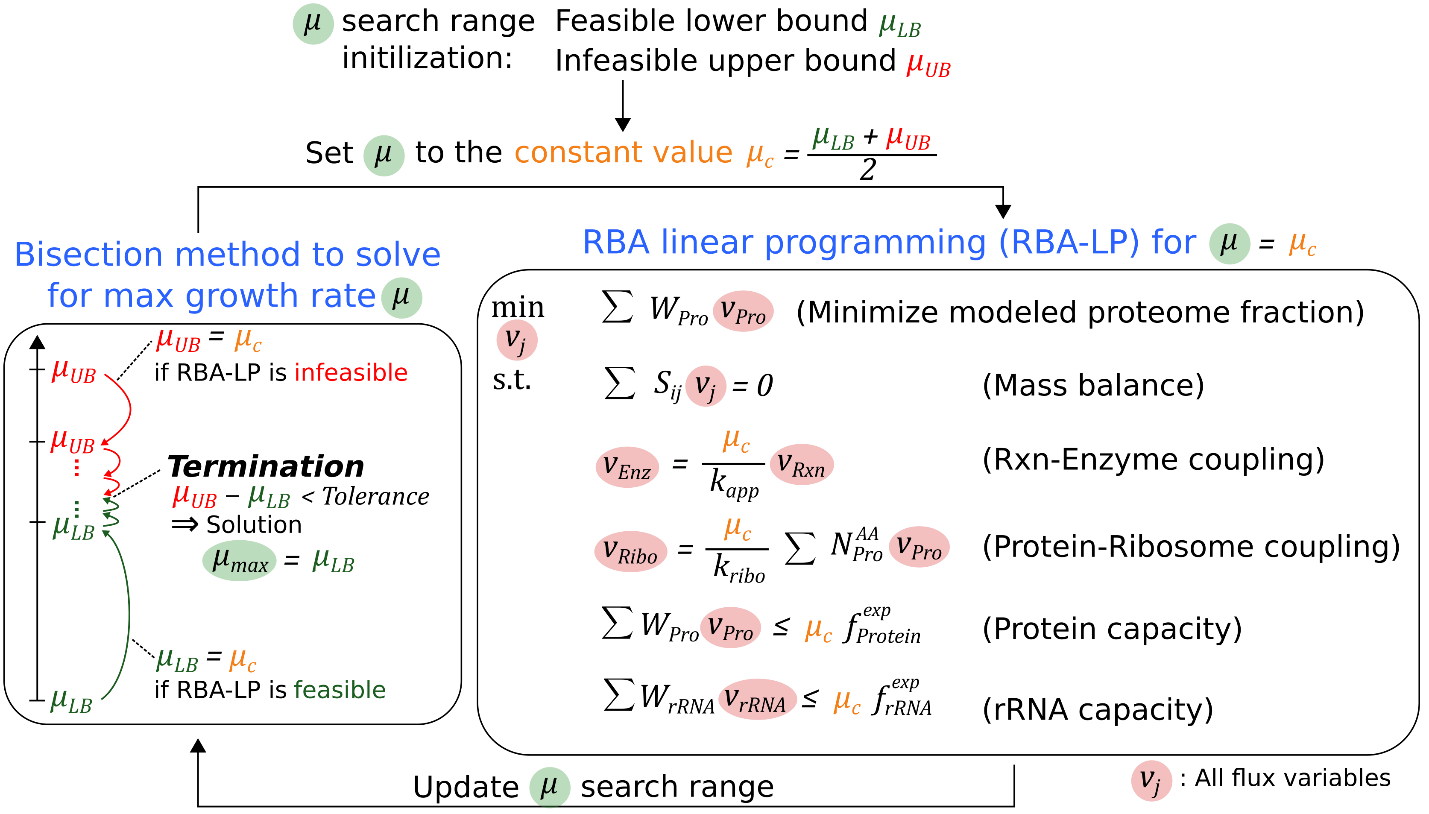

**Figure S1.2**. Overview of mathematical formulation in *sc*RBA.

### A.1. Sets, indexes, and variables

Let $i$ be an element in the set $I$ of all molecular species, which comprises of following subsets: (i) metabolites $I_{M}$, (ii) proteins $I_{P}$, (iii) enzymes $I_{E}$, and (iv) biomass precursors $I_{B}$.

Let $j$ be an element in the set $J$ of all reactions, which comprises of following subsets: (i) reactions in metabolic network $J_{Rxn}$ (including metabolic, transport, and exchange reactions), (ii) protein translation reactions $J_{Psyn}$, and (iii) ribosome synthesis reactions $J_{Ribo}$, (iv) enzyme synthesis reactions $J_{Esyn}$, (v) enzyme load reactions $J_{Load}$, and (vi) biomass synthesis reactions $J_{Bio}$. Variables $v_{j}$ are create on the index $j\in J$. It is to be noticed that all fluxes $j\in J\text{ \textbackslash}J_{Bio}$ have the unit of mmol gDW^-1^ h^-1^, whereas the biomass dilution fluxes $j\in J_{Bio}$ have the unit of g gDW^-1^ h^‑1^. Molar flux is converted to mass flux by applying the molecular weights in g mmol^-1^ as stoichiometric coefficients for the biomass precursors in their synthesis reactions.

In Section A.3 and A.4, we denote the specific growth rate constraint variable as $\mu$ (in h^-1^).

### A.2. Mass balance equations formulation

The mass balance constraints can be written as $\boldsymbol{Sv}=\boldsymbol{0}$ in the matrix-and-vector form, where $\boldsymbol{S}$ is the stoichiometric matrix, and $\boldsymbol{v}$ is the column vector containing all flux variables. The concepts of reaction stoichiometry, stoichiometric matrix, and mass balance equations have been explained in the a primer on flux balance analysis (FBA) by Orth et al., 2010 ^1^. In this document, we apply the fundamentals of FBA and reconstruct the components of the matrix $\boldsymbol{S}$ in RBA. The rows of $\boldsymbol{S}$ are indexed by the set of all molecular species and the columns of $\boldsymbol{S}$ or rows of the column vector $\boldsymbol{v}$ are indexed by the set of all reactions.

Throughout the text, stoichiometric coefficients (at a particular (row, column) position in the matrix $\boldsymbol{S}$) can be positive or negative, indicating that a particular metabolite (indexed by a row of $\boldsymbol{S}$) is being produced or consumed by a particular reaction (indexed by a column of $\boldsymbol{S}$), respectively. For referencing purposes, we define sub-matrices of $\boldsymbol{S}^{\boldsymbol{M}\text{-}\boldsymbol{Rxn}}$ and $\boldsymbol{S}^{\boldsymbol{M}\text{-}\boldsymbol{Psyn}}$ containing coefficients of metabolites in metabolic network and protein/RNA synthesis reactions, respectively. We define sub-matrix $\boldsymbol{S}^{\boldsymbol{P}\text{-}\boldsymbol{Esyn}}$ containing coefficients of protein subunits in enzyme synthesis reaction. We define sub-matrix $\boldsymbol{S}^{\boldsymbol{B}\text{-}\boldsymbol{Bio}}$ containing coefficients of biomass precursors sinking into macromolecule group and subsequently biomass. We also define sub-matrices $\boldsymbol{M}^{\boldsymbol{P}\text{-}\boldsymbol{Psyn}}$ and $\boldsymbol{M}^{\boldsymbol{E}\text{-}\boldsymbol{Esyn}}$ mapping proteins and enzymes to their respective synthesis (respectively), $\boldsymbol{M}^{\boldsymbol{E}\text{-}\boldsymbol{Load}}$ mapping enzymes to enzyme load distribution reactions, and $\boldsymbol{M}^{\boldsymbol{M}\text{-}\boldsymbol{Bio}}$ mapping metabolites to dilution reactions for biomass precursors, respectively. Finally, we define sub-matrices $\boldsymbol{W}^{\boldsymbol{M}}$ and $\boldsymbol{W}^{\boldsymbol{P}}$ containing the molecular weights (in g mmol^-1^) of metabolites and proteins/RNAs, as biomass precursors. Mentioned terms will be defined and explained as we go along. We provide the Table 1 below to show the locations of the sub-matrices in $\boldsymbol{S}$.

**Table 1**. Location of sub-matrices in the stoichiometric matrix $\boldsymbol{S}$

|  |  | Reactions ($J$) | | | | | |
| --- | --- | --- | --- | --- | --- | --- | --- |
|  |  | Metabolic network  ($J_{Rxn}$) | Biomass synthesis  ($J_{Bio}$) | Protein & RNA synthesis ($J_{Psyn}$) | Ribosome synthesis ($J_{Ribo}$) | Enzyme synthesis ($J_{Esyn}$) | Enzyme load distribution $(J_{Load}$) |
| Molecular species ($I$) | Metabolites ($I_{M}$) | $\boldsymbol{S}^{\boldsymbol{M}\text{-}\boldsymbol{Rxn}}$ | $\boldsymbol{M}^{\boldsymbol{M}\text{-}\boldsymbol{Bio}}$ | $\boldsymbol{S}^{\boldsymbol{M}\text{-}\boldsymbol{Psyn}}$ |  |  |  |
|  | Proteins & RNAs ($I_{P}$) |  |  | $\boldsymbol{M}^{\boldsymbol{P}\text{-}\boldsymbol{Psyn}}$ | $\boldsymbol{S}^{\boldsymbol{P}\text{-}\boldsymbol{Ribo}}$ | $\boldsymbol{S}^{\boldsymbol{P}\text{-}\boldsymbol{Esyn}}$ |  |
|  | Enzymes ($I_{E}$) |  |  |  |  | $\boldsymbol{M}^{\boldsymbol{E}\text{-}\boldsymbol{Esyn}}$ | $\boldsymbol{M}^{\boldsymbol{E}\text{-}\boldsymbol{Load}}$ |
|  | Biomass precursors ($I_{B}$) | $\boldsymbol{W}^{\boldsymbol{M}}$ | $\boldsymbol{S}^{\boldsymbol{B}\text{-}\boldsymbol{Bio}}$ | $\boldsymbol{W}^{\boldsymbol{P}}$ |  |  |  |

#### A.2.1. Mass balance equations for metabolites

The general form of mass balance equations for metabolites is:

$$\sum_{j\in J_{Rxn}} S_{ij}^{M\text{-}Rxn}v_{j}\text{ }+\sum_{j\in J_{Psyn}} S_{ij}^{M\text{-}Psyn}v_{j}\text{ }+\sum_{j\in J_{Bio}} S_{ij}^{M\text{-}Bio}v_{j}=0,\text{ }\forall i\in I_{M}$$

where the four terms, from left to right, correspond to net production or consumption flux of molecular species across the metabolic network, protein/RNA synthesis reactions, and biomass synthesis reactions.

Most metabolites are produced and consumed only by reactions in metabolic network ($j\in J_{Rxn}$), thus their mass balance equations are simplified to as in FBA:

$$\sum_{j\in J_{Rxn}} S_{ij}^{M\text{-}Rxn}v_{j}=0$$

Extending from FBA to RBA, some metabolites that are precursors to proteins and RNAs have their stoichiometric coefficients $S_{ij}^{M\text{-}Psyn}$ non-zero for their synthesis reactions (i.e., $j\in J_{Psyn}$). The indexes of proteins & RNAs and their synthesis reactions are explained in Section A.2.2. In addition, some metabolites are also consumed for other biomass constituents via biomass synthesis reactions ($j\in J_{Bio}$). Metabolites and corresponding biomass synthesis reactions are mapped using $M_{ij}^{M\text{-}Bio}$. We will detail how to reconstruct those synthesis reactions sequestering metabolites to establish metabolite mass balance.

##### Metabolites to proteins

There are two types of proteins: modeled and dummy. Modeled proteins are subunits of enzymes and ribosomes whereas dummy protein is used to model the requirement of non-enzymatic and non-ribosomal proteins as well as unused enzymatic proteins. We use the term “dummy protein” to be consistent with published literatures (e.g., in ME-model ^2^ and ETFL-model ^3^). The general form of modeled‑protein translation reaction $j\in J_{Psyn}$ containing only molecular species without stoichiometric coefficients $S_{ij}^{M\text{-}Psyn}$ (which are instead shown in Table 2) is (in which $AA$ denotes amino acid):

$$\sum_{AAs} Charged\text{-}tRNA+\sum_{Cofactors} Cofactor+GTP+ATP+H_{2}O\underset{\to}{}Protein+Biomass\text{-}Protein+\sum_{AAs} Charged\text{-}tRNA+GDP+ADP+P_{i}+H^{+}$$

**Table 2**. Stoichiometric coefficients $S_{ij}^{M\text{-}Psyn}$ in translation reaction of protein of length n.

| **Species** | **Reaction coefficient** | **Source** |
| --- | --- | --- |
| Protein | 1 |  |
| Biomass-Protein | Molecular weight of protein (in g mmol^-1^) | Protein sequence from Saccharomyces Genome Database (SGD) ^4^ |
| Charged-tRNAs^*^ | Number of a specific amino acid molecules in the protein sequence |  |
| Uncharged-tRNAs^*^ | Negative number of a specific amino acid molecules in the protein sequence |  |
| Cofactors | Number of cofactor molecules required for the protein | Uniprot ^5^ |
| GTP | 2n | Translation elongation’s energy requirements ^6,7^ |
| GDP | -2n |  |
| ATP | 1 |  |
| ADP | -1 |  |
| H_2_O | -(2n+1) |  |
| P_i_ | 2n+1 |  |
| H^+^ | 2n+1 |  |

^*^There are 20 charged-tRNAs and 20 uncharged-tRNAs corresponding to 20 amino acids

Charged-tRNA is the carrier of amino acid for protein synthesis and becomes uncharged-tRNA once used. Per amino acid (out of 20), uncharged-tRNA is converted to charged‑tRNA by the following aminoacyl-tRNA synthetase reaction (that is part of the metabolic network, i.e., $j\in J_{Rxn}$):

$$Uncharged\text{-}tRNA+Amino acid+ATP\underset{\to}{} Charged\text{-}tRNA+\text{ }AMP+PP_{i}$$

Lumped in the protein translation reaction is the energy demand in the formed of GTP and ATP hydrolysis reactions (i.e., $ATP\text{/}GTP+H_{2}O\underset{\to}{}ADP\text{/}GDP+P_{i}+H^{+}$).

Protein translation reaction creates two protein molecular species, $Protein$ ($i\in I_{P}$) and $Biomass\text{-}Protein$ ($i\in I_{B}$), feeding into enzyme and biomass synthesis, respectively. This is because we need to evaluate protein as both enzyme/ribosome precursor (i.e., see Section A.4 for coupling constraints) and capacity load consumer (i.e., see Section A.3 for capacity constraints).

In contrast, dummy protein translation reaction creates only $Biomass\text{-}ProteinDummy$ (i.e., different than $Biomass\text{-}Protein$ to track proteome allocation, see Section A.3), because dummy protein does not feed to enzyme/ribosome synthesis. Thus, except for the absence of $Protein$, the generic reaction form and stoichiometric coefficients of the dummy protein are the same with those of modeled proteins. Dummy protein assumes the median length of all proteins in the model (i.e., 401 amino acids) and has amino acid composition matches experimental measurements (i.e., in ^8^).

##### Metabolites to RNAs

There are six specific rRNA species (i.e., 18S, 25S, 5.8S, 5S, 15S, and 21S) and three dummy RNA species (i.e., dummy rRNA, tRNA, and mRNA), indexed as $i\in I_{P}$. The difference is that the specific rRNAs feed into ribosome synthesis reaction whereas dummy RNAs are created to model mRNA and tRNA requirements and unused rRNA. Their synthesis reactions are indexed as $j\in J_{Psyn}$. The general form of their synthesis reactions containing only molecular species without stoichiometric coefficients $S_{ij}^{M\text{-}Psyn}$ (which are instead shown in Table 3) is:

$$ATP+CTP+GTP+UTP\underset{\to}{}rRNA+Biomass\text{-}rRNA+PP_{i}$$

**Table 3.** Stoichiometric coefficients $S_{ij}^{M\text{-}Psyn}$ in synthesis reaction of an rRNA.

| **Metabolite** | **Reaction coefficient** | **Source** |
| --- | --- | --- |
| rRNA | 1 |  |
| Biomass-RNA | Molecular weight of rRNA (in g mmol^-1^) | RNA sequence from RNAcentral database ^9^ |
| ATP | Negative number of a specific nucleotide molecules in the rRNA sequence |  |
| CTP |  |  |
| GTP |  |  |
| UTP |  |  |
| PPi | Length of rRNA sequence |  |

rRNA synthesis reaction produces two copies of rRNA: $rRNA$ ($i\in I_{P}$) and $Biomass\text{-}rRNA$ ($i\in I_{B}$) which participates in ribosome synthesis (and subsequently protein-ribosome coupling constraint) (see Section A.4) and capacity constraint (see Section A.3), respectively. This is because we need to evaluate rRNA as both ribosome precursor (i.e., see Section A.4 for capacity constraints) and capacity load consumer (i.e., see Section A.3 for capacity constraints).

In contrast, because dummy RNA does not feed to ribosome synthesis, dummy RNA synthesis reaction creates only biomass precursor species $i\in I_{B}$, following these reactions (without stoichiometric coefficients):

$$ATP+CTP+GTP+UTP\underset{\to}{}Biomass\text{-}rRNA+PP_{i}$$

$$ATP+CTP+GTP+UTP\underset{\to}{}Biomass\text{-}mRNA+PP_{i}$$

$$ATP+CTP+GTP+UTP\underset{\to}{}Biomass\text{-}tRNA+PP_{i}$$

Since we only need to capture space occupancy in gram per gram dried weight, dummy RNA assumes the length of one and has A/C/G/U composition matches experimental measurements (i.e., as recorded in the GSM *iSace*1144). (Since transcription is not modeled, RNA length is not relevant, unlike translation and protein length)

##### Metabolites to biomass precursors

In our RBA model, instead of modeling metabolic growth requirements using a single biomass reaction (see FBA and biomass reaction papers for more details ^1,10^) whose coefficients indicate the ratio of precursor sequestration, we create individual sink reactions $j\in J_{Bio}$ and impose on them flux values derived from biomass composition. Our configuration to model biomass precursor requirements is detailed in Section A.3. Here, we mentioned only the necessary information to establish the mass balance equation. For a metabolite $i(\text{Bio})\in I_{M}$ that is considered a biomass precursor (see Section A.3), there is a corresponding biomass precursor synthesis reaction $j(\text{Met-to-Bio})\in J_{Bio}$. Let us define the sub‑matrix $\boldsymbol{M}^{\boldsymbol{M}\text{-}\boldsymbol{Bio}}$ encoding the one-to-one mapping relationship of the $(i\left( \text{Bio} \right),\text{ }j(\text{Met-to-Bio}))$ pair. $M_{ij}^{M\text{-}Bio}=-1$ if the pair $(i,j)$ matches the one-to-one mapping and $M_{ij}^{M\text{-}Bio}=0$ otherwise. The general form of biomass precursor sink reactions for a metabolite $i$ with the molecular weight $MW_{i}$ (in g mmol^-1^) is:

$$Metabolite_{i}\underset{\to}{}(MW_{i}) Biomass\text{-}Precursor$$

#### A.1.2. Mass balance equations for proteins and RNAs

The general form of mass balance equations for protein/RNA is:

$$\sum_{j\in J_{Psyn}} M_{ij}^{P}v_{j}\text{ }+\sum_{j\in J_{Esyn}} S_{ij}^{P\text{-}Esyn}v_{j}\text{ }+\sum_{j\in J_{Ribo}} S_{ij}^{P\text{-}Ribo}v_{j}=0,\text{ }\forall i\in I_{P}$$

where the three terms, from left to right, correspond to protein/RNA synthesis flux (i.e., protein synthesis reaction), consumption flux to enzyme synthesis, and consumption flux to ribosome synthesis.

First, indexes of proteins/RNAs and their synthesis reactions in the model needs to be explained. A protein/RNA $i\left( \text{Pro} \right)\in I_{P}$ is produced by a corresponding protein/RNA synthesis reaction $j\left( \text{Pro} \right)\in J_{Psyn}$ (see Section A.2.1 for reaction reconstruction). Let us define the sub‑matrix $\boldsymbol{M}^{\boldsymbol{P}}$ encoding the one-to-one mapping relationship of the $(i\left( \text{Pro} \right),\text{ }j(\text{Pro}))$ pair. $M_{ij}^{P}=1$ if the pair $(i,j)$ matches the one-to-one mapping and $M_{ij}^{P}=0$ otherwise.

On enzyme synthesis, the general form of an enzyme synthesis reaction ($j\in J_{Esyn}$) containing only proteins and enzymes without stoichiometric coefficients is:

$$Proteins\underset{\to}{}Enzyme$$

In an enzyme synthesis reaction, stoichiometric coefficients $S_{ij}^{P\text{-}Esyn}$ of proteins equal to negative values of numbers of protein subunits in enzyme, sourced from the Uniprot database ^5^. Due to an incomplete coverage of all enzymes in the model, missing enzyme stoichiometry is established using Oftadeh et al., 2021’s workflow ^3^. Briefly, enzyme’s subunit stoichiometry is assumed to be the same with its isozyme with subunit information (i.e., obtained from genome-scale metabolic model’s gene-protein-reaction rule), if that applies. Otherwise, enzyme is assumed to be monomeric.

On ribosome synthesis, there are two ribosome synthesis reactions for nucleus (i.e., indexed as $j\left( \text{Ribonuc} \right)$) and mitochondrial (i.e., indexed as $j(\text{Ribomito})$) ribosomes, respectively. This means that $J_{Ribo}=\{j\left( \text{Ribonuc} \right),\text{ }j(\text{Ribomito})\}$. The general form of ribosome synthesis reaction containing only molecular species without stoichiometric coefficients $S_{ij}^{M\text{-}Ribo}$ is:

$$Proteins+rRNAs\underset{\to}{}\emptyset$$

Ribosome is not explicitly modeled as a molecular species as shown in the ribosome synthesis reaction. Under the steady-state mass balance assumption, ribosome synthesis flux strictly equals to the flux of ribosome diluting to daughter cells. Instead of introducing ribosome (as a molecular species) and ribosome sink reaction, mass balance of ribosome is implicitly modeled as shown in the synthesis reaction.

A nucleus ribosome molecule is synthesized from 80 protein and 4 rRNA subunits (rRNAs 18S, 25S, 5.8S, and 5S) ^4^. Their stoichiometric coefficients $S_{ij}^{M\text{-}Ribo}$ are all -1 for the reaction $j\left( \text{Ribonuc} \right)$. Two gene paralogs are present for each of 55 protein subunits (denoted using the suffices “A” and “B” in the standard name on SGD) due to the whole genome duplication event. Since the number of combinations of all possible paralog-specific ribosome is too large, only the version “A” presents in the model to calculate resource allocation. A mitochondrial ribosome molecule is synthesized from 74 protein and 2 rRNA subunits (rRNAs 15S and 21S) ^4,11^. Their stoichiometric coefficients $S_{ij}^{M\text{-}Ribo}$ are all -1 for the reaction $j(\text{Ribomito})$.

#### A.1.3. Enzyme load distribution reactions of total enzyme pool

The general form of mass balance equations for enzymes ($\forall i\in I_{E}$) is:

$$\sum_{j\in J_{Esyn}} M_{ij}^{E}v_{j}+\sum_{j\in J_{Load}} M_{ij}^{E\text{-}Load}v_{j}=0,\text{ }\forall i\in I_{E}$$

where the two terms, from left to right, correspond to enzyme production flux from protein precursors and enzyme consumption to enzyme load (and subsequently to biomass production).

An enzyme $i\left( \text{Enz} \right)$ is produced by a corresponding enzyme synthesis reaction $j\left( \text{Enz} \right)\in J_{Esyn}$ (see Section A.2.2 for reaction reconstruction). Let us define the sub-matrix $\boldsymbol{M}^{\boldsymbol{E}}$ encoding the one-to-one mapping relationship of the $(i\left( \text{Enz} \right),\text{ }j(\text{Enz}))$ pair. $M_{ij}^{E}=1$ if the pair $(i,j)$ matches the one-to-one mapping and $M_{ij}^{E}=0$ otherwise.

An enzymatic reaction in the model is catalyzed by an enzyme load, which is a partition of whole enzyme pool. This is because an enzyme can catalyze multiple reactions or the forward and reverse direction of a reaction. Enzyme load distribution reactions are used in RBA model to map the oftentimes enzyme-reaction mappings that are not one-to-one. In RBA model, reversible reactions are also split to irreversible forward and reverse versions. In addition, the same enzymatic reactions catalyzed by isozymes are also split into components catalyzed by respective isozymes. To illustrate, we provide examples on reconstruction of enzymatic reactions and enzyme load distribution reactions in RBA model. In those examples, an irreversible reaction $Rxn1$ is equivalent to $Rxn1F$ and a reversible reaction $Rxn1$ is split into $Rxn1F$ and $Rxn1R$, where suffices $F$ and $R$ indicate forward and reverse reactions, respectively.

**Example 1**: A single enzyme ($EnzA$) catalyzing a single irreversible reaction ($Rxn1$).

| **Enzyme** | **Enzyme load** | **Reaction** |
| --- | --- | --- |
| $EnzA$ | $LoadA1F$ | $Rxn1F(LoadA1F)$ |

**Example 2**: A single enzyme ($EnzA$) catalyzing a single reversible reaction ($Rxn1$).

| **Enzyme** | **Enzyme load** | **Reaction** |
| --- | --- | --- |
| $EnzA$ | $LoadA1F$ | $Rxn1F(LoadA1F)$ |
|  | $LoadA1R$ | $Rxn1R(LoadA1R)$ |

**Example 3**: Two isozymes ($EnzA$ and $EnzB$) catalyzing a single reversible reaction ($Rxn1$). Here, the two versions of $Rxn1F$ are considered two separate reactions.

| **Enzyme** | **Enzyme load** | **Reaction** |
| --- | --- | --- |
| $EnzA$ | $LoadA1F$ | $Rxn1F(LoadA1F)$ |
| $EnzB$ | $LoadB1F$ | $Rxn1F(LoadB1F)$ |

**Example 4**: A single enzyme ($EnzA$) catalyzing two irreversible reactions ($Rxn1$ and $Rxn2$).

| **Enzyme** | **Enzyme load** | **Reaction** |
| --- | --- | --- |
| $EnzA$ | $LoadA1F$ | $Rxn1F(LoadA1F)$ |
|  | $LoadA2F$ | $Rxn2F(LoadA2F)$ |

**Example 5**: Two isoenzymes ($EnzA$ and $EnzB$) catalyzing two irreversible reactions ($Rxn1$ and $Rxn2$).

| **Enzyme** | **Enzyme load** | **Reaction** |
| --- | --- | --- |
| $EnzA$ | $LoadA1F$ | $Rxn1F(LoadA1F)$ |
|  | $LoadA2F$ | $Rxn2F(LoadA2F)$ |
| $EnzB$ | $LoadB1F$ | $Rxn1F(LoadB1F)$ |
|  | $LoadB2F$ | $Rxn2F(LoadB2F)$ |

Overall, enzyme-to-load mapping can be one-to-one or one-to-many and load-to-reaction mapping is strictly one-to-one. Now, back to mass balance equations involving total enzyme pools and enzyme loads, let us define enzyme load distribution reaction $j\left( \text{Load} \right)\in J_{Load}$. The general form of an enzyme load distribution reaction is:

$$Enzyme\underset{\to}{}\emptyset$$

Enzyme load, which is essentially enzyme, is not explicitly modeled as a molecular species. Under the steady-state mass balance assumption, enzyme production strictly equals to the flux of enzyme diluting to daughter cells. Instead of introducing enzyme load (as a molecular species) and enzyme sink reaction, mass balance of enzyme load is implicitly modeled as shown in the synthesis reaction.

We define the sub-matrix $\boldsymbol{M}^{\boldsymbol{E}\text{-}\boldsymbol{Load}}$ encoding the mapping relationship between an enzyme $i\left( \text{Enz} \right)\in I_{E}$ and enzyme load distribution reactions $j(\text{Load})$. $M_{ij}^{E\text{-}Load}=-1$ if the pair $(i,j)$ matches a recorded enzyme-to-load relationship and $M_{ij}^{E\text{-}Load}=0$ otherwise.

#### A.1.4. Mass balance of biomass precursors

Through Section A.2.1 and A.2.2, we introduce biomass precursors ($i\in I_{B}$) coming from metabolic network and protein/RNA synthesis reactions. Those initial precursors are further spooled into their respective macromolecule group and then sink reactions are create for the group. The general form of mass balance equation for biomass precursor is:

$$\sum_{j\in J_{Rxn}} W_{ij}^{M}v_{j}\text{ }+\sum_{j\in J_{Psyn}} W_{ij}^{P}v_{j}\text{ }+\sum_{j\in J_{Bio}} S_{ij}^{B\text{-}Bio}v_{j}=0,\text{ }\forall i\in I_{B}$$

where $\boldsymbol{W}^{\boldsymbol{M}}$ and $\boldsymbol{W}^{\boldsymbol{P}}$ are the molecular weight matrices. $W_{ij}^{M}=MW_{i}$ ($i\in I_{M}$) and $W_{ij}^{P}=MW_{i}$ ($i\in I_{P}$) if the index $i$ of biomass precursor matches its synthesis reaction $j$. Otherwise, $W_{ij}^{M}=0$ and $W_{ij}^{P}=0$. Here, $MW_{i}$ is the molecular weight of metabolite for $i\in I_{M}$ and of protein/RNA for $i\in I_{P}$. Stoichiometric coefficients $S_{ij}^{B\text{-}Bio}$ represent the biomass precursor synthesis network shown in Figure S.1.3. We set up biomass synthesis network to incorporate the *S. cerevisiae*’s measurements of biomass composition that changes with growth rate ^8^. In addition, we also account for proteome pool allocation toward non-enzymatic and non-ribosomal proteins, which is 45% ^3^, and account for RNA pool allocation toward tRNA and mRNA, which is 15% and 5% ^12^, respectively.

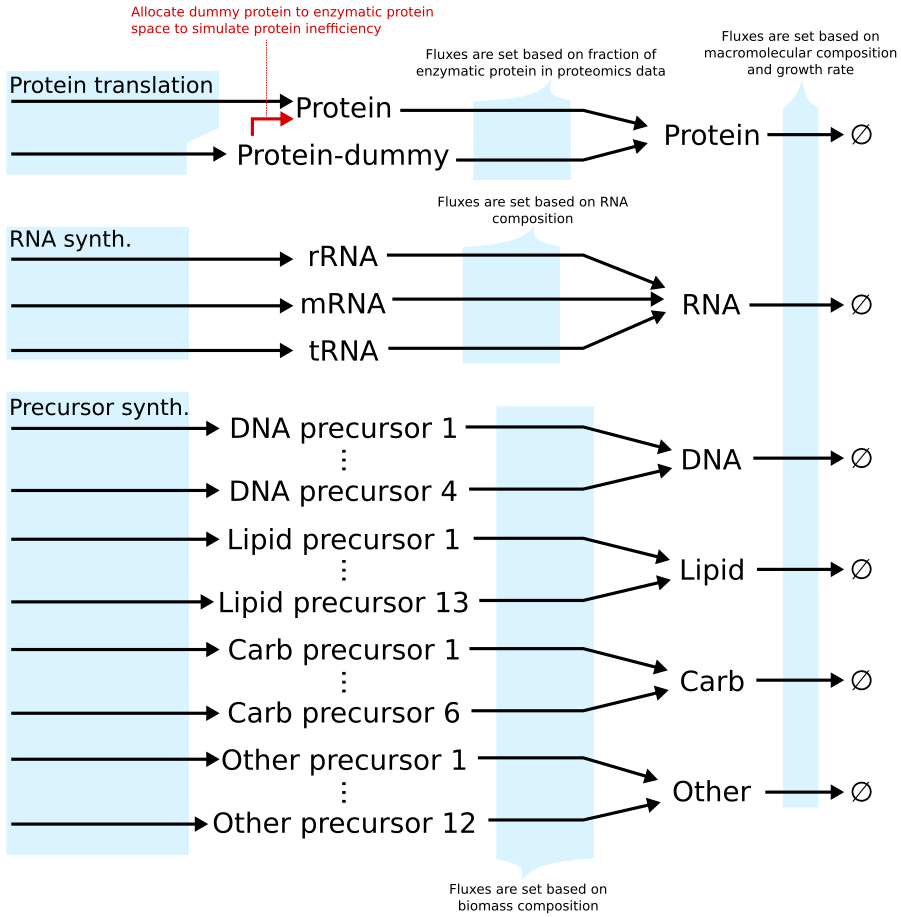

**Figure S1.3**. Biomass precursor synthesis network. The goal of the network is to facilitate the implementation of capacity constraints and to incorporate biomass composition measurements.

### A.3. Biomass production and capacity constraints formulation

There are six constraints that impose flux value. Their general form is as follows:

$$v_{j}=\mu f_{j}^{Bio}, j\in\left\{ Protein,RNA,DNA,Lipid,Carb,Other \right\}\subset J_{Bio}$$

The flux imposition indicates that the amount of biomass precursor production ($v_{j}$, in g gDW^-1^ h^‑1^) equals to the amount diluting to the daughter cells, which is the specific growth rate ($\mu$, in h^-1^) times the mass fraction in biomass (g gDW^-1^). Based on biomass composition measurements ^8,13^, protein, RNA, and carbohydrate fractions in biomass varies with growth rate whereas the other fractions remain relatively constant. We thus approximate those fractions using a linear fit to the available experimental composition data points for protein fraction (from $\mu\in[0.022, 0.4]$, R^2^ = 0.67) and RNA fraction (from $\mu\in\left[ 0.022,0.211 \right]$, R^2^ = 0.72). Carbohydrate fraction is determined by subtracting protein, RNA, and other fractions. The parameters are provided in the following Table 4:

**Table 4**. Mass fraction parameter value (in g gDW^-1^) or equation

| **Mass fraction parameter** | **Value or equation** |
| --- | --- |
| Protein | $f_{Protein}^{Bio}=0.3736+0.3292 \mu$ |
| RNA | $f_{RNA}^{Bio}=0.0402+0.1370 \mu$ |
| Carb | $f_{Carb}^{Bio}=0.4693-0.4662 \mu$ |
| Lipid | $f_{Lipid}^{Bio}=0.0697$ |
| DNA | $f_{DNA}^{Bio}=0.0039$ |
| Metal (part of other) | $f_{Metal}^{Bio}=0.0254$ |
| Cofactor (part of other) | $f_{Cofactor}^{Bio}=0.0048$ |
| Sulphate (part of other) | $f_{SO4}^{Bio}=0.0030$ |
| Phosphate (part of other) | $f_{Pi}^{Bio}=0.0101$ |

To account for the proteome allocation towards non-enzymatic and non-ribosomal proteins (i.e., modeled as dummy protein), we added the following flux impositions:

$$v_{Protein\text{-}Enz\&Ribo}=\mu f_{Enz\&Ribo}^{Protein} f_{Protein}^{Bio}$$

$$v_{Protein\text{-}Dummy}=\mu f_{Dummy}^{Protein} f_{Protein}^{Bio}$$

where $f_{Enz\&Ribo}^{Protein}=0.55$ and $f_{Dummy}^{Protein}=0.45$ are the experimentally observed mass fraction of protein allocated to enzymatic & ribosomal protein and other proteins, respectively ^3^.

To account for the RNA allocation towards mRNA and tRNA, we added the following flux impositions:

$$v_{rRNA}=\mu f_{rRNA}^{RNA} f_{RNA}^{Bio}$$

$$v_{tRNA}=\mu f_{tRNA}^{RNA} f_{RNA}^{Bio}$$

$$v_{mRNA}=\mu f_{mRNA}^{RNA} f_{RNA}^{Bio}$$

where $f_{rRNA}^{RNA}=0.8$, $f_{tRNA}^{RNA}=0.15$, $f_{mRNA}^{RNA}=0.05$ are the experimentally observed mass fractions of rRNA, tRNA, and mRNA in total RNA pool, respectively.

Because mass balance constraint enforces the protein & RNA production fluxes to be equal to the protein & RNA sink (see Section A.2.2), and flux constraints impose flux values to the protein & RNA sink fluxes (see Section A.2.4 and this section), the protein and rRNA capacity constraints are implicitly modeled. The equivalent explicit protein and rRNA capacity constraints, respectively, are:

$$v_{Protein-Enz\&Ribo}=\sum_{j\in J_{Proteins}\subset J_{Psyn}} MW_{i} M_{ij}^{P} v_{j}=\mu f_{Enz\&Ribo}^{Protein} f_{Protein}^{Bio}$$

$$v_{rRNA}=\sum_{j\in J_{rRNA}\subset J_{Psyn}} MW_{i} M_{ij}^{P} v_{j}=\mu f_{rRNA}^{RNA} f_{RNA}^{Bio}$$

where $J_{Proteins}(\subset J_{Psyn}$) is the set of protein translation flux (which excludes r/t/mRNA synthesis fluxes), and $J_{rRNA}(\subset J_{Psyn})$ is the set of rRNA synthesis flux (which excludes protein translation fluxes and t/mRNA synthesis fluxes). A protein translation reaction creates two copies: $Protein$ and $Biomass\text{-}Protein$, thus the flux imposition on $Biomass\text{-}Protein$ sink restrict the protein translation flux and subsequently how much enzyme and ribosome can be made from $Protein$. The same applies for rRNA synthesis flux creating $rRNA$ and $Biomass\text{-}rRNA$.

### A.4. Coupling constraints formulation

#### A.4.1. Reaction-Enzyme coupling constraints

The general form of reaction-enzyme coupling constraints is:

$$\mu v_{j(\text{Rxn})}\leq k_{app}\left( j\left( \text{Rxn} \right) \right) M_{j\left( \text{Rxn} \right),j\left( \text{Load} \right)} v_{j\left( \text{Load} \right)},\text{ } j(\text{Rxn})\in J_{Rxn}$$

where $v_{j(\text{Rxn})}$ is the flux of enzyme catalyzed reaction $j\left( \text{Rxn} \right)\in J_{Rxn}$, $k_{app}(j(\text{Rxn}))$ is the apparent kinetic turnover number (unit of h^-1^) of enzyme on reaction $j(\text{Rxn})$, $v_{j(\text{Load})}$ is the enzyme load synthesis flux $j\left( \text{Load} \right)\in J_{Load}$, and $M_{j\left( \text{Rxn} \right),j\left( \text{Load} \right)}$ is the matrix mapping the one-to-one relationship of reaction and enzyme load (see Section A.2.3). $M_{j\left( \text{Rxn} \right),j\left( \text{Load} \right)}=1$ if ($j\left( \text{Rxn} \right),j(\text{Load})$) is the matching pair and $M_{j\left( \text{Rxn} \right),j\left( \text{Load} \right)}=0$ otherwise.

To derive the reaction-enzyme coupling constraint, we use the non-indexed variables and parameters for simplicity. Reaction-Enzyme coupling constraints are modeled with the first-order kinetic rate law with respect to enzyme concentration (i.e., $[Enz]$), $v_{j(\text{Rxn})}\leq k_{app}\left[ Enz \right]$, where $k_{app}$ is the apparent kinetic turnover number parameter. Multiplying that inequality by $\mu$ yields $\mu v_{j(\text{Rxn})}\leq k_{app}v_{j(\text{Enz})}$, where the enzyme production rate (i.e., $v_{j(\text{Enz})}$) equals to the rate of enzyme diluting to daughter cells (i.e., $\mu[Enz]$) in steady-state mass balance of enzymes. This is because as the cell maintains the concentration $[Enz]$ and grows at the rate of $\mu$, the enzyme dilution rate is $\mu[Enz]$. To model enzyme catalyzing multiple reactions, enzyme loads partitioned from total enzyme pool are used instead (by replacing $v_{j(\text{Enz})}$ with $v_{j(\text{Load})}$), yielding $\mu v_{j(\text{Rxn})}\leq k_{app}v_{j(\text{Load})}$.

#### A.4.2. Protein-Ribosome coupling constraints

The protein-ribosome coupling constraints for ribosomal ($j\left( \text{Ribonuc} \right)\in J_{Ribo}$) and mitochondrial ($j\left( \text{Ribomito} \right)\in J_{Ribo}$) ribosomes are:

$$\mu\sum_{j\in J_{Pnuc}} N_{j}^{AA}v_{j}\text{ }\leq\text{ }k_{Ribo}v_{j(\text{Ribonuc})}$$

$$\mu\sum_{j\in J_{Pmito}} N_{j}^{AA}v_{j}\text{ }\leq\text{ }k_{Ribo}v_{j(\text{Ribomito})}$$

where $N_{j}^{AA}$ is the number of amino acids in the sequence of the protein being produced by the protein translation reaction $j\in J_{Psyn}$ (see Section A.1.2 for protein indexing), $k_{Ribo}$ is the ribosome efficiency (unit of amino acids per ribosome per second for referrals in text, amino acids per ribosome per hour in implementation for unit consistency within the model), $J_{Pnuc}\subset J_{Psyn}$ is the set of proteins translated by nucleus ribosome, and $J_{Pmito}$ is the set of proteins translated by mitochondrial ribosome. Ribosome efficiency $k_{ribo}$ is 13.2 amino acids per ribosome per second (x3600 for per hour). The number is derived from the literature-reported value 10.5 ^14^. We found the updated value by increase from 10.5 incrementally by 0.1 until the simulation where growth rate is fixed to the highest experimental value of 0.49 h^-1^ ^15^ is feasible. Based on mitochondrial genome annotation ^16^, the eight proteins in the model being translated by the mitochondrial ribosome are Q0045, Q0080, Q0085, Q0105, Q0130, Q0140, Q0250, and Q0275. The rest of the proteins are translated by the nucleus ribosome.

To derive the protein-ribosome coupling constraint, we use the non-indexed variables and parameters for simplicity. The total amount of proteins is limited by the following protein-ribosome coupling constraint of $\sum_{Proteins} N_{Pro}^{AA} v_{Pro}\leq k_{Ribo}[Ribo]$, where $N_{Pro}^{AA}$ is the length of protein sequence, $v_{Pro}$ is the protein translation flux, $k_{Ribo}$ is the ribosome efficiency, and $[Ribo]$ is the ribosome concentration. Protein-Ribosome coupling constraint indicates that to perform protein translation at a certain rate the cell needs to maintain a corresponding ribosome concentration. Since ribosome dilution rate (i.e., $\mu[Ribo]$) equals to production rate (i.e., $v_{Ribo}$), multiplying that equation by $\mu$ yields $\mu\sum_{Proteins} N_{Pro}^{AA} v_{Pro}\leq k_{Ribo}v_{Ribo}$.

### B. Software implementation

Software is available at <https://github.com/maranasgroup/scRBA>. The scripts are currently set up for reconstruction and simulation of *S. cerevisiae* metabolism, but they can be adapted for other organisms with sufficient molecular biology knowledge and mathematical modeling and programming skill. General instructions for reconstruction and simulation will be described below.

#### B.1. Input files requirements

The following data are collected and stored in excel spreadsheets format (.xlsx), except for the genome-scale metabolic model that is stored in COBRApy JSON format (.json) ^17^.

| **Input file** | **Description** |
| --- | --- |
| **Common path**: ./scRBA/build_model/input/  (the actual file names are different on the github, but the same (capitalized) prefix tags are used) | |
| GSM.json | Genome-scale model file in COBRApy format |
| BIOMASS.xlsx | Biomass composition |
| PROTEIN.xlsx | Protein precursors stoichiometry and molecular weights |
| PROTEIN_dummy.xlsx | Parameters on dummy protein |
| ENZYME.xlsx | Enzyme precursors stoichiometry and mappings to reactions |
| RIBOSOME_nucleus.xlsx | Nucleus ribosome composition |
| RIBOSOME_mitochondrial.xlsx | Mitochondrial ribosome composition |
| RNA.xlsx | RNA compositions |
| PARAMS.xlsx | Apparent turnover numbers for enzymes |

#### B.2. Model reconstruction

Go to the following directory: ./scRBA/build_model/

**1) Execute the script**: A01_build_excel_stoich_for_GAMS.ipynb

This will yield the excel spreadsheet with all reactions compiled:

./model/RBA_stoichiometry.xlsx

**2) Execute the script**: A02_build_GAMS_Sij_and_fluxBounds.ipynb

This will yield the following GAMS model files below. The user needs to manually copy (or write their own scripts to copy) these files to: ./scRBA/GAMS/model/. This was done to prevent overwriting a GAMS file in-working by GAMS file newly created by the python scripts (which could be faulty).

| **GAMS file** | **Description** |
| --- | --- |
| **Common path**:  Files are created in ./scRBA/build_model/model/  Files are manually copied to and called by GAMS at this path: ./scRBA/GAMS/model/ | |
| RBA_species.txt | List of all (macro)molecules |
| RBA_rxns.txt | List of all reactions |
| RBA_sij.txt | Stoichiometric matrix coefficients |
| RBA_rxns_EXREV.txt | A reaction subset. List of nutrient uptake reactions (exchange, reverse). All of them will be turned off by GAMS script to prevent uptake of nutrients that are not in the media. Uptake of nutrients in the media has to be (over)written through “RBA_rxns_EXREV_<media_id>.txt”. |
| RBA_rxns_EXFWD.txt | A reaction subset. List of byproduct secretion reactions (exchange, forward). (Created but not used in the GAMS script. Don’t worry if you cannot figure out the purpose of this file.) |
| RBA_rxns_rxnmetabolicnetwork.txt | A reaction subset. List of reactions in the original metabolic network of GSM. (Used in parameterization, to calculate minimal metabolic flux distribution, i.e., minimizing sum of fluxes) |
| RBA_rxns_enzsyn.txt | A reaction subset. List of synthesis reaction for total enzyme pool. (Used in parameterization, to calculate enzyme concentration from protein concentration) |
| RBA_rxns_enzload.txt | A reaction subset. List of synthesis reaction for individual enzyme loads constituting total enzyme pool. (Used in parameterization, to identify which reactions can be active based on proteomics data). |
| RBA_rxns_prowaste.txt | A reaction subset. List of protein waste reactions (which sink the protein to void). These sink reactions are created to debug model protein synthesis and are turned off by default. (Turning these reactions on will create a lot of degrees of freedom to the optimization and significantly inflate the run time) |

The following GAMS model files encoding the nutrient availability in the media has to be written manually by the user:

| **GAMS file** | **Description** |
| --- | --- |
| **Common path**: ./scRBA/GAMS/model/ | |
| RBA_rxns_EXREV_mineralMinimum.txt | All uptake reactions of minerals |
| RBA_rxns_EXREV_YNB.txt | All uptake reactions of minerals and vitamins in YNB media |
| RBA_rxns_EXREV_YP.txt | All uptake reactions of minerals, vitamins in YNB media, and supplemented amino acids |

**3) Execute the script**: A03_build_GAMS_RBA_constraints.ipynb

Ribosome-Protein coupling constraints are coded directly in runRBA.gms. Thus, this script only writes the following files:

| **GAMS file** | **Description** |
| --- | --- |
| **Common path**:  Files are created in ./scRBA/build_model/model/  Files are manually copied to and called by GAMS at this path: ./scRBA/GAMS/model/ | |
| RBA_enzCapacityConstraints_declares.txt | Declared indexes of reaction-enzyme coupling constraints |
| RBA_enzCapacityConstraints_eqns.txt | Inequality reaction-enzyme coupling constraints. This file is created but is not further used because running inequality constraints take significantly longer time that would prevent the model from being practical. |
| RBA_enzCapacityConstraints_eqns_equality_version.txt | Equality reaction-enzyme coupling constraints (being used). Equality constraint implementation is practical (reduce solving time), valid, and consistent with the RBA-LP objective, which is minimizing protein capacity usage. |
| RBA_rxns_prosyn.txt | A reaction subset. List of protein translation reactions. These reactions are called by ribosome-protein coupling constraints. |
| RBA_proteinLength.txt | Protein sequence lengths. These parameters are called by ribosome-protein coupling constraints. |

**4) k_app_ parameters**

k_app_ parameters are stored in ./scRBA/input/ (with the prefix “RBA_kapp_”) and are called by reaction-enzyme coupling constraints. The k_app_ needs to be in per hour for the input files. See parameterization methods section for how they are estimated for *sc*RBA model.

#### B.3. Model simulation

Here, only general script structure and guidance are provided since different setup could be involved for different analysis. (Example) Software scripts for *sc*RBA simulation creating the results shown in the manuscript is available at <https://github.com/maranasgroup/scRBA>.

Generic GAMS files are available at ./scRBA/GAMS/. Case studies simulation are recommended to be organized to a different folder. Specific GAMS script needs to be copied to that case study folder and execute within there. Every GAMS script .gms file is accompanied by a GAMS_settings .txt file that contains a path link to the files called by the .gms script.

Individual RBA-LP optimization is run by runRBA.gms through the command line, with growth rate specified (see below for example run for growth rate of 0.1 h^-1^).

gams runRBA.gms --mu=0.1 o=/dev/null

Binary search to iterative run RBA-LP is executed through binary_search.py with run settings specified in binary_search_options.py (generic example file is stored in ./scRBA/pycore/binary_search/).

Raw outputs are processed by python functions (stored in ./scRBA/pycore/) to yield the organized results in JSON format (see examples in <https://github.com/maranasgroup/scRBA>).

The paths to the following GAMS scripts are available for case studies mentioned in the manuscript:

- Flux variability:
  - GAMS core scripts: ./scRBA/GAMS/vary_flux/
  - Python exec scripts: ./scRBA/vary_flux/
- Max production analysis:
  - GAMS core scripts: ./scRBA/GAMS/application/
  - Python exec scripts: ./scRBA/application/

#### B.4. Parameterization method and software implementation

**Methods**

k_app_ parameterization workflow is illustrated in Figure S.1.4. To estimate k_app_, we first estimated enzyme concentrations (i.e., in mmol gDW^-1^) from experimental protein concentrations (i.e., in mmol gDW^-1^). For an enzyme composed of a single protein, its concentration is equal to protein concentration divided by protein subunit stoichiometric coefficient (e.g., 1 for monomer, 2 for homodimer). For an enzyme composed of different proteins, its molar concentration is equal to the least abundant protein subunit. Mathematically, it is the minimum of values, each of which is calculated by dividing protein molar concentration to its enzyme subunit stoichiometric coefficient. In our calculations, a few protein subunits, even for essential enzymes such as ATP synthase, were not measured and thus inference of enzyme concentration relies on the available protein subunit measurements. After that, we calculated intracellular fluxes from experimental extracellular fluxes in a two-step procedure. First, we determine essential (measurement “gap‑filled”) enzyme that was not recorded in experimental data by minimizing the sum of flux of reactions catalyzed by enzymes that are not measured. Then, we determine the flux distribution from an optimization procedure wherein sum of metabolic fluxes catalyzed by measured and “gap-filled” enzymes are minimized. At the end, k_app_ value is calculated by dividing the metabolic flux by the enzyme concentration. For cases where multiple reactions are catalyzed by the same enzyme or multiple isozymes catalyze the same reaction, since we cannot ascertain individual enzyme-reaction catalysis load, we assume the same k_app_ value for all reactions and enzymes involved and the sum of metabolic fluxes and the sum of enzyme concentration values are used in k_app_ calculation.

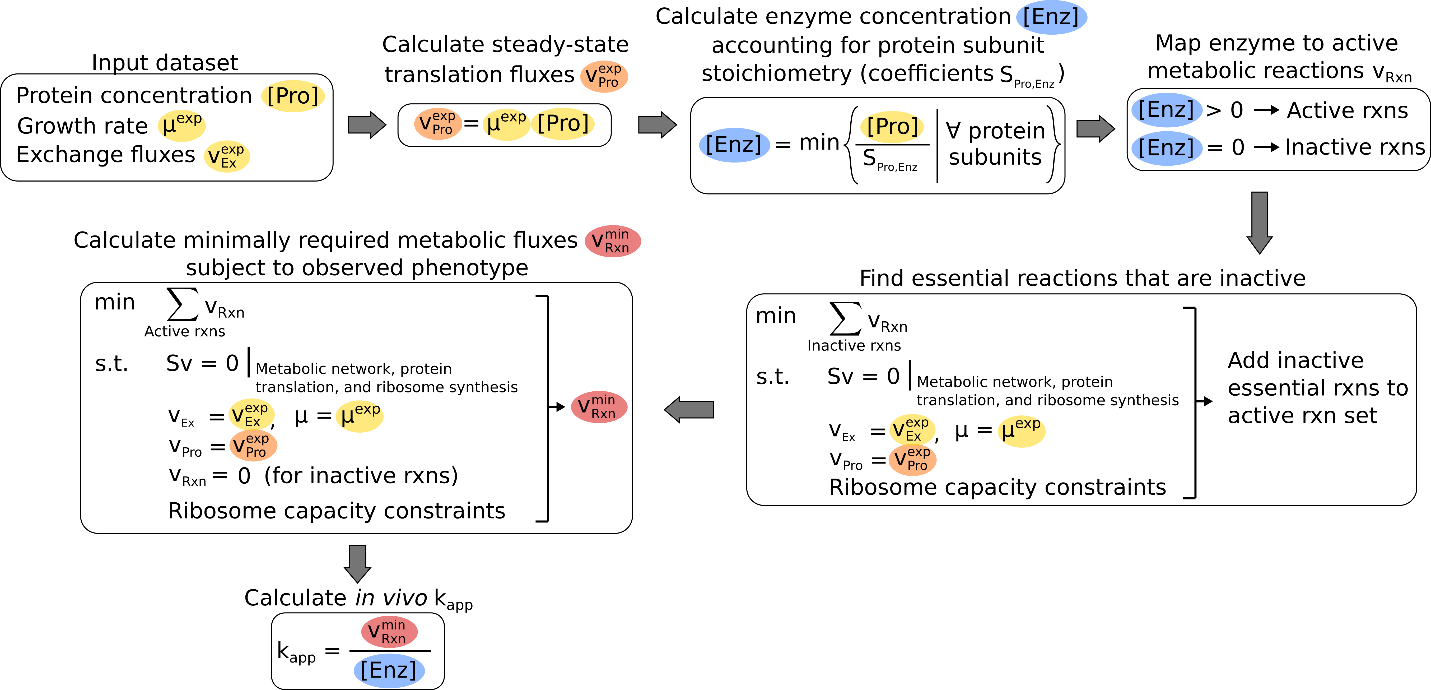

**Figure S.1.4**. *In vivo* k_app_ parameterization workflow. Exchange metabolic fluxes and protein concentrations data were used as the inputs to calculate k_app_ from intracellular metabolic fluxes and enzyme concentrations, accounting for metabolic reaction and protein subunit stoichiometry.

**Software implementation**

Proteomics data needs to be manually processed by users. Scripts are only meant to be used as examples since they are specific to *S. cerevisiae*. After proteomics data processing, then:

Step 1: Estimate enzyme concentrations from protein concentrations, accounting for protein subunit stoichiometry

- GAMS core: ./scRBA/GAMS/parameterization/enz_from_proteome/
- Python exec

./scRBA/parameterization/kapp/datasets/<id>/B1_enz_from_proteome.py

Step 2: Rescue reactions from inactive set and put into active set. This is to account for the fact that proteomics data might be incomplete and miss out enzymes performing essential metabolic functions.

- GAMS core: ./scRBA/GAMS/parameterization/min_flux_violation/
- Python exec

./scRBA/parameterization/kapp/datasets/<id>/B2_min_flux_violation.py

Step 3: Estimate minimal metabolic flux distributions (only allow reactions in active set to carry fluxes) (analogous to parsimonious FBA ^18^)

- GAMS core: ./scRBA/GAMS/parameterization/min_flux_sum/
- Python exec

./scRBA/parameterization/kapp/datasets/<id>/B3_min_flux_sum.py

Step 4: Estimate k_app_ from metabolic flux and enzyme concentration.

- Python exec:

./scRBA/parameterization/kapp/datasets/<id>/C1_calculate_kapp.ipynb

#### B.5. Solver settings

The solver settings need to be recorded in the .opt (text) file and has to be in the same directory as the GAMS script .gms file based on GAMS programming syntax. SoPlex solver was used to solve the RBA-LP linear programming problem, thus the file name is “soplex.opt”. (See examples in <https://github.com/maranasgroup/scRBA>)
