## Supplementary Figures for "Evaluating proteome allocation of *Saccharomyces cerevisiae* phenotypes with resource balance analysis"

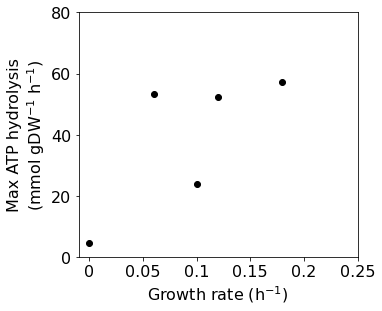

**Supplementary Fig. 1 | Maximal ATP maintenance vs. growth rates under phosphate limited conditions.**
